# Resolving the evolutionary history of bighorn sheep to inform future management: an answer to the California bighorn lineage question

**DOI:** 10.1101/2025.03.13.643091

**Authors:** Joshua P. Jahner, Thomas L. Parchman, Marjorie D. Matocq, Mike Cox, Rachel S. Crowhurst, Lanie M. Galland, Shelby M. Burdo, Michael R. Buchalski, Joshua M. Hallas, Soraia Barbosa, David W. Coltman, Samuel Deakin, Holly B. Ernest, Sierra M. Love Stowell, Hollie Miyasaki, Kevin L. Monteith, Annette Roug, Helen Schwantje, Robert S. Spaan, Thomas R. Stephenson, Jace Taylor, Lisette P. Waits, John D. Wehausen, Clinton W. Epps

## Abstract

Although translocations can be effective for augmenting and restoring wild populations, they can disrupt native patterns of genetic structure, diversity, and local adaptation, thereby hampering conservation efforts. Managers must weigh potential costs and benefits of choosing well-differentiated donor individuals that could confer a boost to genetic diversity while avoiding outbreeding depression or ecological mismatch. This decision is more daunting when taxonomy is unclear or debated. For example, bighorn sheep (*Ovis canadensis*) populations in the United States that have been managed as the “California” lineage (part of the formerly recognized subspecies *O. c. californiana*) originate from serial translocations sourced from populations in British Columbia, resulting in reduced genetic diversity and elevated risk of inbreeding. After research on skull shape and RFLP analysis of mtDNA failed to find support for that subspecies, some jurisdictions treated the California lineage as part of the Rocky Mountain subspecies (*O. c. canadensis*) and mixed individuals in subsequent translocations, in part to increase genetic diversity of bottlenecked populations. Yet detailed genetic data addressing validity of those putative lineages were lacking. We reconstructed the genetic history of bighorn sheep by sampling the major putative subspecies or lineages, focusing on native (remnant) genetic variation, and generating high-throughput DNA sequencing data (∼15,000-25,000 SNPs). Complementary phylogenetic and population genetic analyses supported the distinctiveness of four bighorn lineages at levels corresponding to subspecies. Our results confirm the genetic identity of the no longer putative California bighorn lineage, answering a question that puzzled geneticists and managers for decades. Moving forward, we recommend that managers 1) maintain the natural variation held in native populations by protecting them from intentional translocations or unintentional mixing with nearby populations; 2) prioritize within-lineage translocations for population augmentation or repatriation to previously occupied regions; and 3) cautiously consider any translocations that would lead to mixing of distinct evolutionary lineages.

## Introduction

Identifying the evolutionary history of populations helps establish conservation priorities and guide management decisions (Hendry *et al*., 2010). Populations that have been separated over deep evolutionary time frames are expected to be genetically and phenotypically diverged as a result of both isolation and adaptation to different conditions, and may exhibit some degree of reproductive isolation (Ord & Summers, 2015). Restoration efforts in response to local or regional extirpation of a species typically aim to limit mixing among divergent lineages to minimize the risk of outbreeding depression and maladaptation (IUCN/SSC, 2013). This can be particularly challenging when cryptic or previously-unrecognized levels of differentiation exist (e.g., giraffes; Winter *et al*., 2018). Yet, the magnitude of evolutionary divergence below the species level is often poorly understood and have been infrequently explored with modern genomics approaches. As a result, translocation efforts aimed at restoring biodiversity may inadvertently lead to mixing of differentially adapted lineages or those that have evolved some degree of reproductive isolation (Garcia de Leaniz *et al*., 2007; Wiedmann & Sargeant, 2014; Malaney *et al*., 2015). Because wildlife management decisions are strongly driven by taxonomic delineations, it is imperative that taxonomy properly reflects our understanding of evolutionary divergence. Even though reliable and defensible criteria for distinguishing taxonomic categories below the species level remain elusive and problematic (Fraser & Bernatchez, 2001; Turbek *et al*., 2023; Clavero *et al*., 2024), characterizing the evolutionary history of populations allows managers to make informed decisions concerning the maintenance of distinct evolutionary lineages and the ecological contexts in which they will be conserved.

In North America, large mammals, particularly herbivores, have been subject to extensive conservation management, and translocations have often been used to restore populations following widespread extirpation and range collapse in the 19th and early 20th centuries (Kallman, 1987). Translocation has played a central role in restoration of species such as white-tailed deer (*Odocoileus virginianus*), elk (*Cervus canadensis*), pronghorn (*Antilocapra americana*), mountain goats (*Oreamnos americanus*), and bighorn sheep (*Ovis canadensis*) (e.g., Gann *et al*., 2020; Chafin *et al*., 2021; Whiting *et al*., 2023; Sacks *et al*., 2024). In some cases, translocations relied on the best available information regarding evolutionary history; in other cases, little regard was paid to such concerns (Wiedmann & Sargeant, 2014; Wild Sheep Working Group, 2015). Bighorn sheep are an iconic big game species of western North America with ecological and cultural significance. They are found in rugged reaches of western North America, and exemplify managed species that now exist as a patchwork of native remnants and reestablished populations. They occupy habitats spanning a wide range of environmental conditions, from the high alpine of the Rocky Mountains and Sierra Nevada to the hot deserts of Mexico and the southwestern United States. In the late 1800s and early 1900s, bighorn sheep were subjected to overhunting, competition from livestock, and probably most significantly, diseases transmitted from domestic sheep that caused population declines and extirpations throughout much of the historic range (Buechner, 1960; Wehausen *et al*., 2011; Cassirer *et al*., 2018). Over the past 80 years, state, provincial, federal, and tribal partners worked together to reintroduce bighorn sheep to previously occupied habitats (Wild Sheep Working Group, 2015), resulting in widespread recovery across the former range. However, in the course of that restoration, few populations retained fully native genetic ancestry (referred to hereafter as native populations [Epps *et al*., 2010] or remnant populations [Jahner *et al*., 2019]), either because few or no native populations remained in some regions, or because remaining populations were augmented by translocated individuals from geographically distant populations. In some jurisdictions, translocations used nearby source stock that was likely adapted to similar conditions (e.g., California; Bleich *et al*., 2021), yet in others (e.g., Oregon, North Dakota, northern Nevada), source stock came from very distant and different habitats (Singer *et al*., 2000; Whiting *et al*., 2023). Many reintroduction programs followed the subspecies taxonomy proposed by Cowan (1940), which has since been recognized as inadequate in some respects (Wehausen & Ramey, 2000). Although Cowan‘s (1940) analysis was largely based on univariate cranial morphological measurements, subsequent analyses applied genetic and multivariate morphological data (Ramey, 1993; Wehausen & Ramey, 2000) to address differentiation among putative lineages, or explored relationships solely with genetic data (Malaney *et al*., 2015; Buchalski *et al*., 2016; Barbosa *et al*., 2021).

Genetic research on bighorn sheep has a rich history, but most studies have been limited in geographic scope, addressed questions at shallow evolutionary timescales, or used low resolution methods of describing variation. Past studies investigating genetic differentiation have characterized genetic structure among populations and lineages (Ramey, 1993, 1995; Buchalski *et al*., 2015, 2016; Malaney *et al*., 2015; Deakin *et al*., 2020; Love Stowell *et al*., 2020), demonstrated the genetic consequences of translocations (Miller *et al*., 2012; Olson *et al*., 2012; Malaney *et al*., 2015; Gille *et al*., 2019; Jahner *et al*., 2019; Flesch *et al*., 2020), identified major barriers to gene flow (Epps *et al*., 2005, 2018), and quantified the relationships between genetic diversity and isolation in the context of vulnerability to climate change (Creech *et al*., 2020), among other efforts reviewed in Epps *et al*. (2026). This work has influenced the management of bighorn sheep over the past two decades, but many questions regarding historical diversification and evolutionary relationships among putative lineages remain, driven in part by varying resolution of different types of data and analyses. Analyses of whole-genome sequencing data suggest that bighorn sheep diverged from the more northerly distributed thinhorn sheep (*O. dalli*) approximately 2.5 – 2.2 Mya (Santos *et al*., 2021). Systematic evaluation of bighorn sheep has been limited. Ramey (1993, 1995) examined mitochondrial DNA restriction fragment length polymorphisms (RFLP) variation and concluded that within bighorn, Rocky Mountain and California bighorn sheep from western British Columbia formed a clade, whereas desert and southern (Sierra Nevada) populations of California bighorn sheep formed another, with long separation between those clades, refuting Geist‘s (1971) hypothesis that Rocky Mountain and desert bighorn derived from a single southern refugium. Luikart & Allendorf (1996) examined mitochondrial DNA RFLP variation in Rocky Mountain bighorn and found little or no phylogeographic structure. The fossil record suggests that bighorn sheep occupied at least two major refugia throughout the late Pleistocene, including a northern refugium in present day Wyoming established ∼100 kya (Martin & Gilbert, 1978; Wang, 1988) and a southern refugium in the Mojave Desert established ∼300 kya (Geist, 1985). Potentially consistent with that pattern, Buchalski *et al*. (2016) analyzed mitochondrial control region sequences and identified two well-supported clades representing Rocky Mountain (*O. c. canadensis*) and Sierra Nevada bighorn sheep (*O. c. sierrae*), but a third clade consisting of desert bighorn (*O. c. nelsoni*) was not statistically supported. Due to sampling limitations, that dataset was unable to provide further insight into the evolutionary origins of the putative California bighorn sheep lineage (*O. c. californiana*).

Despite the extensive history of bighorn sheep genetic research, uncertainty regarding the evolutionary distinctiveness (or lack thereof) of California bighorn sheep remains a critical knowledge gap that acts as a major impediment to the successful implementation of future management across the northern part of the range. Cowan (1940) described three northern subspecies: California bighorn sheep in western British Columbia (B. C.) and extending south as far as the Sierra Nevada Mountains in California; Audubon’s bighorn sheep (*O. c. auduboni*, now extinct) in eastern Montana and Wyoming, North and South Dakota, and western Nebraska; and Rocky Mountain bighorn sheep, *O. c. canadensis*, extending from eastern B. C. and Alberta south to New Mexico. More statistically sophisticated analyses based on larger sample sizes than those used by Cowan (1940) found that Rocky Mountain, northern populations of California (from B. C. and Washington), and Audubon’s bighorn sheep did not differ significantly in cranial morphology (Wehausen & Ramey, 2000). Cowan’s definition of California bighorn extended that subspecies south through the Cascade Mountains of Washington and Oregon and into the Sierra Nevada of California, with areas of uncertainty indicated in eastern Washington, central Oregon, and northwestern Nevada where the transition to Rocky Mountain bighorn was unclear (Cowan, 1940). On the basis of morphometric cranial analyses and mtDNA RFLP data, Wehausen *et al*. (2005) separated Sierra Nevada populations as a subspecies, *O. c. sierrae*, first proposed by Grinnell (1912). Wehausen & Ramey (2000) also noted that bighorn skulls from extirpated populations in Oregon had desert bighorn morphological characteristics. Native bighorn in Oregon, Washington, and northern Nevada were all extirpated by the early 20th century. Translocations to reestablish bighorn in those regions were partly informed by Cowan’s subspecies designations, and largely used bighorn from western B. C., the only non-Sierra Nevada portion of the range of the California subspecies where native bighorn remained. On the eastern edges of those regions, Rocky Mountain bighorn were used as translocation stock, although southwestern Idaho received California bighorn. A recent genetic investigation of bighorn sheep currently existing in Idaho, Washington, and B. C. found strong differentiation between populations of California and Rocky Mountain bighorn sheep ancestry (Barbosa *et al*., 2021). However, except for most of those from B. C., the study populations all stemmed from translocations, which may have uncertain histories. Moreover, the microsatellite markers used in that study are not ideal for revealing deeper evolutionary history.

Some jurisdictions still manage California bighorn sheep originating from western B. C. as a distinct entity (e.g., Oregon, Nevada), whereas others now treat them as Rocky Mountain bighorn sheep (e.g., Utah). This distinction is important for management due to the very low levels of genetic diversity found in California bighorn sheep populations in the United States, all of which were reintroduced via serial translocations of founders from a few populations in B. C. (e.g., Olson *et al*., 2013). If the two putative lineages are not differentiated from one another, California populations could be augmented with individuals from more genetically diverse Rocky Mountain populations (one such translocation in North Dakota resulted in higher recruitment; Wiedmann & Sargeant, 2014). Alternatively, if California and Rocky Mountain bighorn sheep represent independent evolutionary lineages, translocation of one lineage into the range of the other could have unintended consequences including erosion of native genetic variation, outbreeding depression, and maladaptation.

Here we fill a key knowledge gap in the evolutionary history and future management of bighorn sheep by finally providing an answer to the question of whether the putative California bighorn sheep lineage represents a distinct lineage. We generated a genotyping-by-sequencing (GBS) dataset consisting of remnant bighorn sheep populations, or in some instances translocated populations known to stem from those remnants, from the major putative lineages found in the United States and Canada. Our goals were to: 1) examine patterns of diversification for extant bighorn sheep and identify major lineages within the species, and 2) consider implications of those findings for management and future translocation programs. We expected that extant populations of bighorn sheep recognized as desert, Rocky Mountain (sensu Cowan, 1940), and Sierra Nevada (Wehausen *et al*., 2005) bighorn subspecies would represent well-supported and deeply divergent lineages. We considered two alternate hypotheses for the western B. C. (and translocated descendants) populations of bighorn sheep previously described as California bighorn by Cowan (1940), and subsequently synonymized by Wehausen & Ramey (2000) with Rocky Mountain bighorn: 1) bighorn sheep from the southern Rocky Mountains, north into the Canadian Rockies, and west into the westernmost bighorn sheep populations in British Columbia represent a single evolutionary lineage, exhibiting clinal variation and isolation by distance; or 2) populations in western British Columbia and their translocated descendants, described by Cowan (1940) as part of the California subspecies of bighorn, represent a clearly differentiated evolutionary lineage distinct from all other populations of Rocky Mountain bighorn sheep. Although a full taxonomic revision of bighorn sheep is beyond the scope of this paper, we review where our genetic analyses concur and disagree with the standing taxonomy.

## Methods

### Population sampling and data generation

Twenty populations were selected based on known genetic information, management history, and sample availability (Fig. 1; Table 1). Populations, described by some jurisdictions as herds, are geographically and demographically distinct from other bighorn populations due to the patchy nature of bighorn habitat and previous extirpations, although they are not necessarily isolated from gene flow. Populations were sampled from the three currently recognized subspecies found in Canada and the United States (desert, Rocky Mountain [not including western B. C. populations synonymized by Wehausen & Ramey, 2000], Sierra Nevada), as well as the putative California lineage still extant in western B. C. and their translocated descendants. When possible, multiple populations were selected within the range of each putative subspecies, based on previously published genetic studies (Buchalski *et al*., 2016; Jahner *et al*., 2019; Deakin *et al*., 2020; Barbosa *et al*., 2021). Because our primary objective was to identify historical (i.e., pre-1800s) patterns of genetic structure, potential populations were prioritized if they were thought to have remnant genetic ancestry because they either survived the die-offs of the late 1800s and early 1900s or were later restored with individuals from remnant populations. Of the twenty included populations, eight are known to have received individuals from other sources, including four California populations (Hart Mountain, Kamloops Lake, Sinlahekin, South Okanagan), three Rocky Mountain populations (Antelope Island, Georgetown, Ram Mountain), and the lone Sierra Nevada population (Mount Langley) (Table S1; Wild Sheep Working Group, 2015); of these, Kamloops Lake represents the only likely instance of mixed lineages (Rocky Mountain bighorn were translocated into a nearby location in the 20th century). Many of the samples (blood, tissue, or previously extracted DNA) acquired for this study were originally used in other genetic studies with more restricted geographic and taxonomic scopes (e.g., Jahner *et al*., 2019; Deakin *et al*., 2020; Love Stowell *et al*., 2020; Barbosa *et al*., 2021).

**Table 1:**
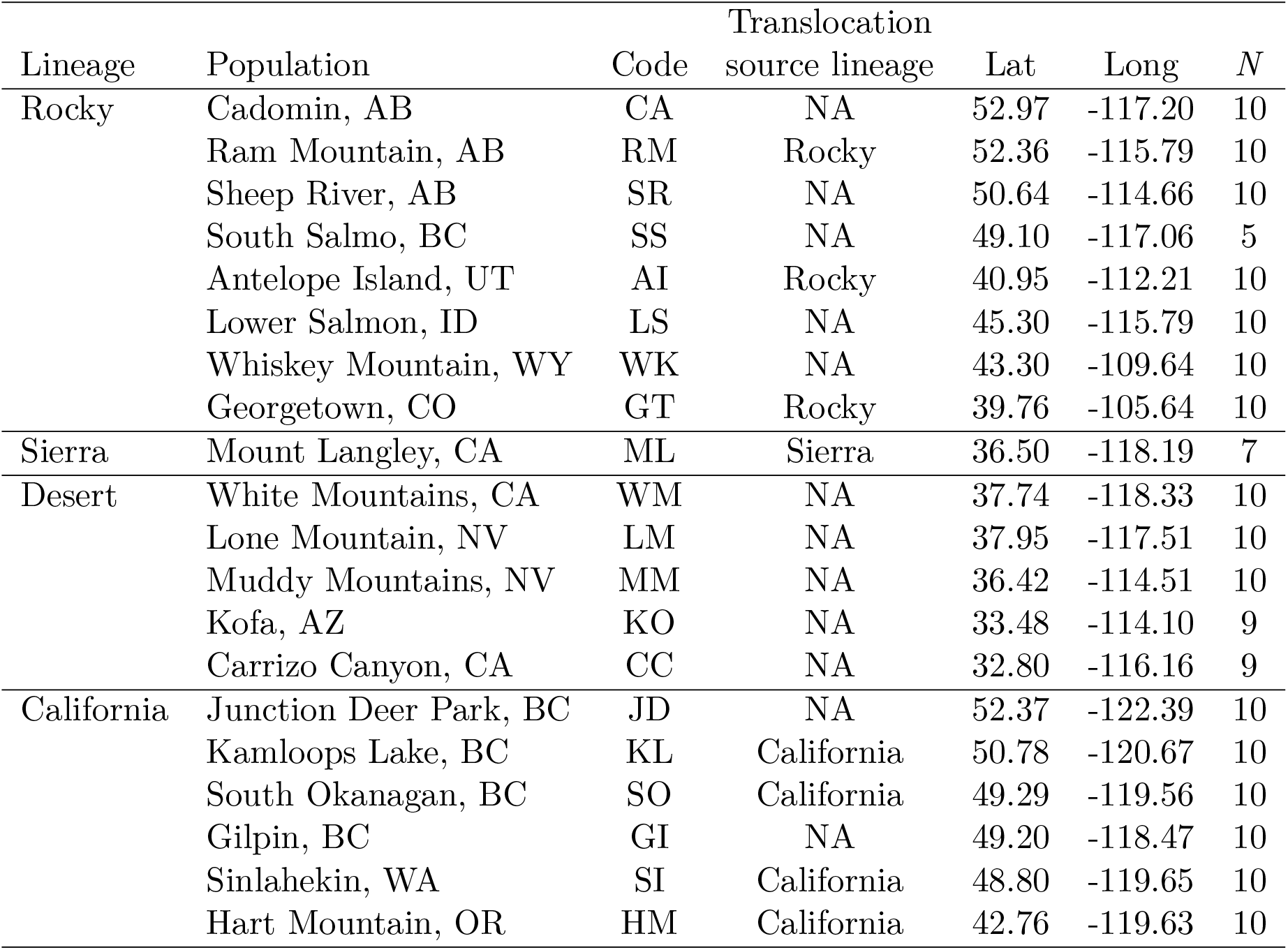
Population information for the 20 bighorn sheep populations included in analyses. Within lineages, populations are roughly ordered by decreasing latitude. Translocations involving populations in this manuscript only occurred using source stock from within the putative lineage. See Table S1 for a full translocation history for the populations included in this study.

**Figure 1:**
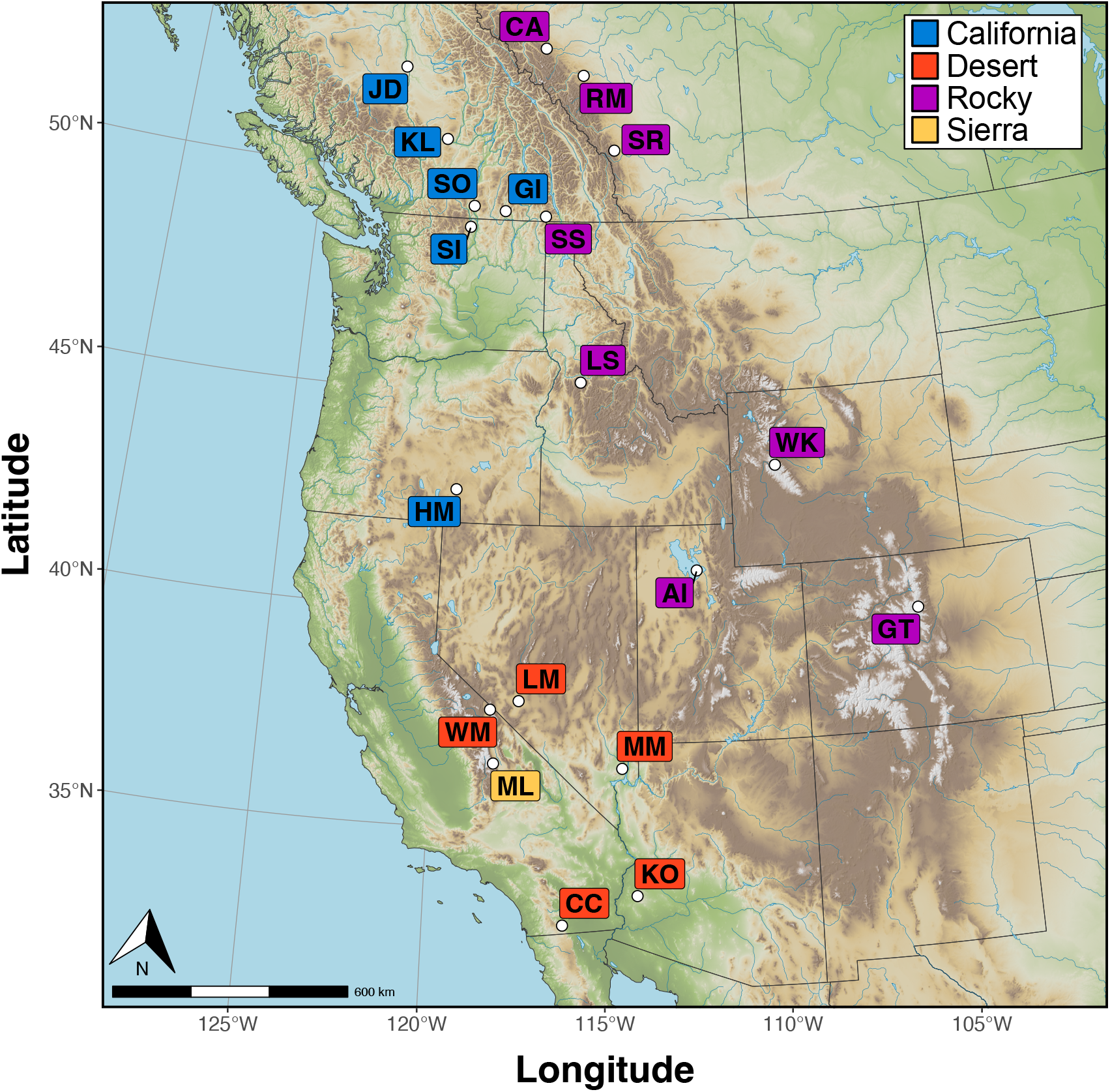
A map of bighorn sheep populations that were included in analyses. See Table 1 for additional population information.

DNA was extracted from samples using DNeasy Blood and Tissue kits (Qiagen Inc.). Sequencing libraries were constructed following the same genotyping-by-sequencing protocol (Parchman *et al*., 2012) used by Jahner *et al*. (2019). DNA was cut using two restriction enzymes (*Eco*RI, *Mse*I), barcoded adaptors unique to each individual were ligated to each DNA fragment, and fragments were amplified with PCR. To reduce the fraction of genome space for sequencing, libraries were size-selected for fragments ranging from 350-450 base pairs (bp) in length using a Pippin Prep (Sage Science). The individuals in this study were part of a larger sequencing effort that included 2,823 individuals sequenced across 20 Illumina HiSeq 2500 lanes (University of Wisconsin-Madison Biotechnology Center; Madison, WI) and two Illumina NovaSeq 6000 lanes with S2 chemistry (University of Texas Genomic Sequencing and Analysis Facility; Austin, TX).

DNA sequences were first filtered of contaminants (e.g., *E. coli*, PhiX) using the tapioca pipeline (https://github.com/ncgr/tapioca) and `bowtie2_db` (Langmead & Salzberg, 2012). Reads were parsed using individual barcodes and split into separate individual fastq files using custom Perl scripts. Individual fastq files are available at Dryad ([dataset] Jahner *et al*., 2025). All reads were aligned to the Rocky Mountain bighorn sheep chromosome-level genome assembly (GCA_001039535.1; Miller *et al*., 2015) using the *aln* and *samse* algorithms of `bwa v0.7.8` (Li & Durbin, 2009), specifying a maximum number of mismatch bases (-n) to four. We also explored analyses using the new unpublished Rocky Mountain bighorn sheep genome (GCF_042477335.2) and found broadly consistent results. Sequence alignment/map (sam) files were converted to binary format (bam) and variant sites were called using `bcftools v1.9` and `samtools v1.9` (Li *et al*., 2009), setting minimum base, mapping, and site qualities of 20 and a minimum genotype quality of 10.

Variant calling was performed separately for population genetic (5 – 10 individuals per population) and phylogenetic (2 individuals per population) analyses using `vcftools v0.1.16` (Danecek *et al*., 2011). A single Stone’s sheep (*O. dalli stonei*) individual was included as an outgroup for phylogenetic analyses. For the population genetic subset, we filtered variant sites with minor allele frequencies (MAF) < 0.03, as well as sites with zero reads for more than 30% of individuals. Phylogenetic trees constructed from reduced representation sequencing data have higher support when including more loci, even with higher proportions of missing data (Wagner *et al*., 2013). As such, we used less stringent filtering for the phylogenetic subset, filtering loci with MAF < 0.02 and sites with zero reads for more than 40% of individuals. To avoid potential genotyping error from over-assembly of paralogous regions, sites with mean depth > 14X per individual or with *F*_*IS*_ *<* −0.5 were also excluded in both subsets.

### Population genetic and phylogenetic analyses

We characterized patterns of genetic ancestry within and between putative lineages using the hierarchical Bayesian model `entropy v2.0` (Gompert *et al*., 2014; Shastry *et al*., 2021). That model uses variation in genotype likelihoods to estimate genotype probabilities for each individual at each locus, as well as ancestry coefficients (*q*) for a specified number of ancestral clusters (K). Importantly, the use of genotype likelihoods (as opposed to unambiguous genotype calls) allows the model to account for inherent uncertainty that arises due to sequencing errors and low sequencing depths. This uncertainty is propogated into genotype probabilities, ancestry coefficients, and downstream population genetic analyses. Five replicate `entropy` chains were run for each of K = 1–10 ancestral clusters, specifying 250,000 MCMC iterations, a 50,000 iteration burnin, and a thinning interval of 40 iterations. After documenting differentiation among the lineages associated with subspecific designations, we quantified potential hierarchical variation within major groups with separate `entropy` models using only California, desert plus Sierra Nevada, and Rocky Mountain subsets, specifying K = 1–5 ancestry clusters, 100,000 iterations, a 50,000 iteration burnin, and a thinning interval of 10 iterations for each. Chain mixing was evaluated with effective sample sizes (ESS) and convergence was assessed with potential scale reduction factors (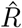; Gelman & Rubin, 1992). Genetic differentiation among and within lineages was further characterized using principal component analysis (PCA) in `R v4.2.1` (R Core Team, 2022) based on the genotype probabilities generated by `entropy`. As above for `entropy` analyses, we conducted PCA on the full set of individuals, as well as separately for each of the three analytical groups (California, desert plus Sierra Nevada, and Rocky Mountain). A neighbor-joining tree was constructed based on pairwise *F* _ST_ (Hudson *et al*., 1992) using the `ape v5.7.1` package (Paradis & Schliep, 2019) in R, with population allele frequencies quantified from genotype probabilities. Finally, we calculated expected heterozygosity (*H*_E_) based on expectations from Hardy-Weinberg equilibrium (i.e., *H*_E_ = 2*pq*).

We inferred phylogenetic trees based on the data subset containing 2 individuals per population. First, the vcf file was converted to Phylip format using `vcf2phylip v2.0` (https://github.com/edgardomortiz/vcf2phylip). Phylogenetic inference was performed using `IQ-TREE v1.6.12` (Nguyen *et al*., 2015), specifying *model finder plus* to find the most appropriate substitution model (Kalyaanamoorthy *et al*., 2017) and applying ultrafast bootstrap with 1,000 replicates (Hoang *et al*., 2018). Based on simulated data, the ultrafast bootstrap is unbiased when support values are higher than 70%, meaning that a value of 90 has a 90% chance of being correct (Minh *et al*., 2013). The tree was rooted with the *O. dalli stonei* individual (not displayed) and visualized using the `ggtree v3.4.4` package (Yu *et al*., 2017) in R.

## Results

After mapping and quality filtering, we retained a population genetic dataset consisting of 190 individuals, 14,622 loci, and a mean depth of 4.42X, and a final phylogenetic dataset consisting of 41 individuals, 24,795 loci and a mean depth of 4.36X. The population genetic data set recovered consistent evidence for hierarchical genetic structure. Overall, `entropy` chains had adequate sampling (effective sample sizes typically greater than 200) and convergence (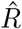 typically close to 1). While the `entropy` model with three genetic ancestries (K) had the lowest DIC (Table S2), models up to K = 8 revealed biologically informative structure where at least one population had individual ancestry coefficients (*q*) of nearly one for each ancestry (Fig. 2). At K = 2, the California bighorn populations split from all other populations, although some Rocky Mountain ancestry was evident in the Kamloops (KL) and South Okanagan (SO) populations. At K = 3, the Rocky Mountain populations were delineated from the desert and Sierra Nevada populations, although three of the four southernmost Rocky Mountain populations (Georgetown, GT; Lower Salmon, LS; Whiskey Mountain, WK) had split ancestry coefficients, suggestive of additional structure at a higher K (Fig. 3). Indeed, these three populations formed their own ancestry group at K ≥ 5. The translocated Antelope Island (AI) population separated as its own ancestry at K ≥ 4. The Sierra Nevada population (Mount Langley, ML) and the geographically proximate White Mountains (WM) population were differentiated from the other desert populations at K = 6 and from each other at K = 7. At K = 8, a north to south ancestry cline emerged for the remaining desert populations (Lone Mountain, LM to Muddy Mountains, MM to Kofa, KO to Carrizo Canyon, CC). Additional `entropy` analyses based on three data subsets (only California, desert plus Sierra Nevada, and only Rocky Mountain) were broadly consistent with the results from the full dataset (Table S3; Figs. S1, S2, S3). PCA produced results qualitatively similar to those described above for `entropy`, with differentiation evident among the three major groups (California, desert plus Sierra Nevada, and Rocky Mountains (Fig. 4). These analyses also illustrated differentiation among most sampled populations within each lineage that followed regional patterns (Fig. 4).

**Figure 2:**
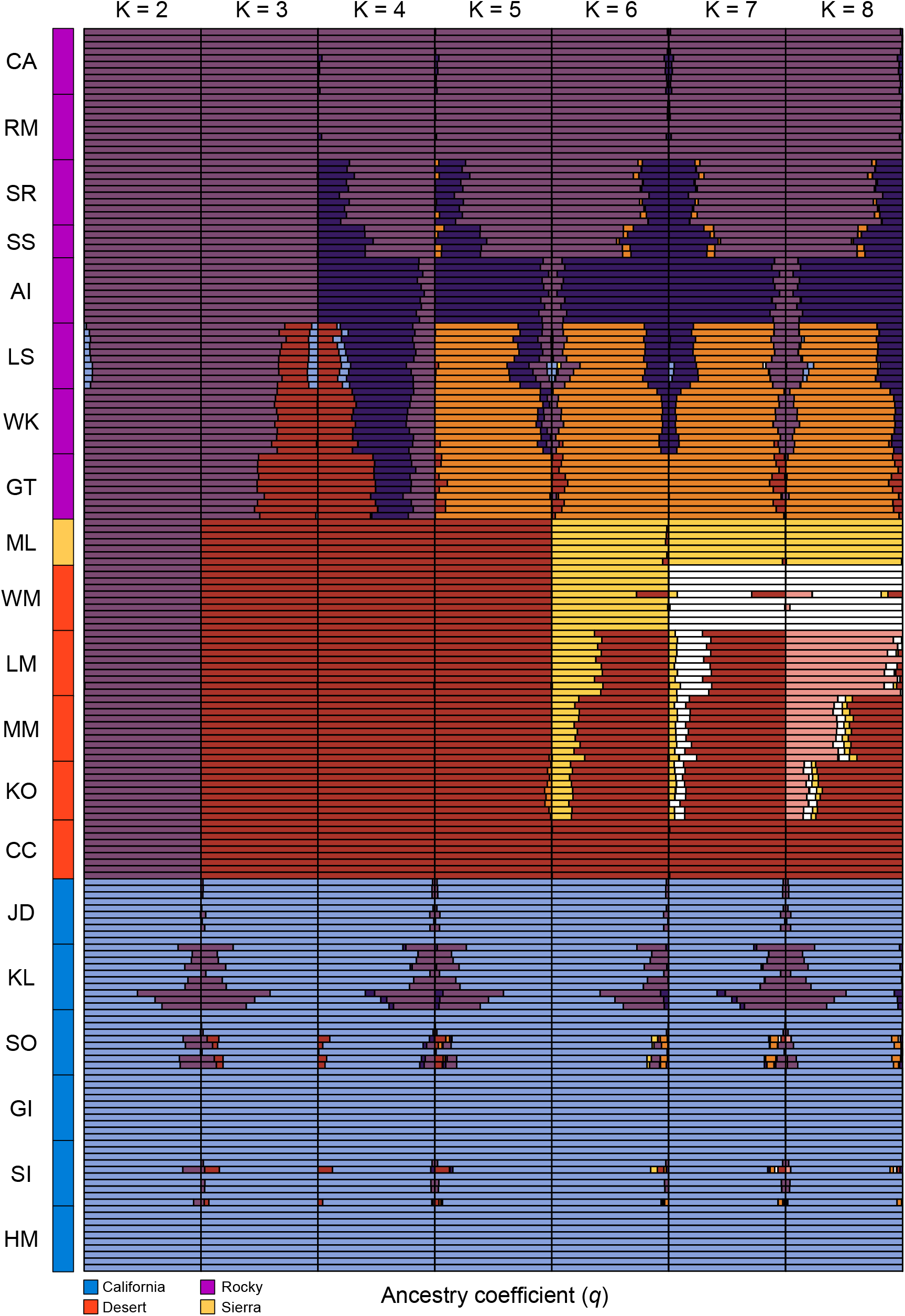
Individual ancestry coefficients (*q*) are shown for full dataset `entropy` models ranging from 2 – 8 genetic clusters (K). Each row represents an individual bighorn sheep, and individuals are sorted by population and lineage.

**Figure 3:**
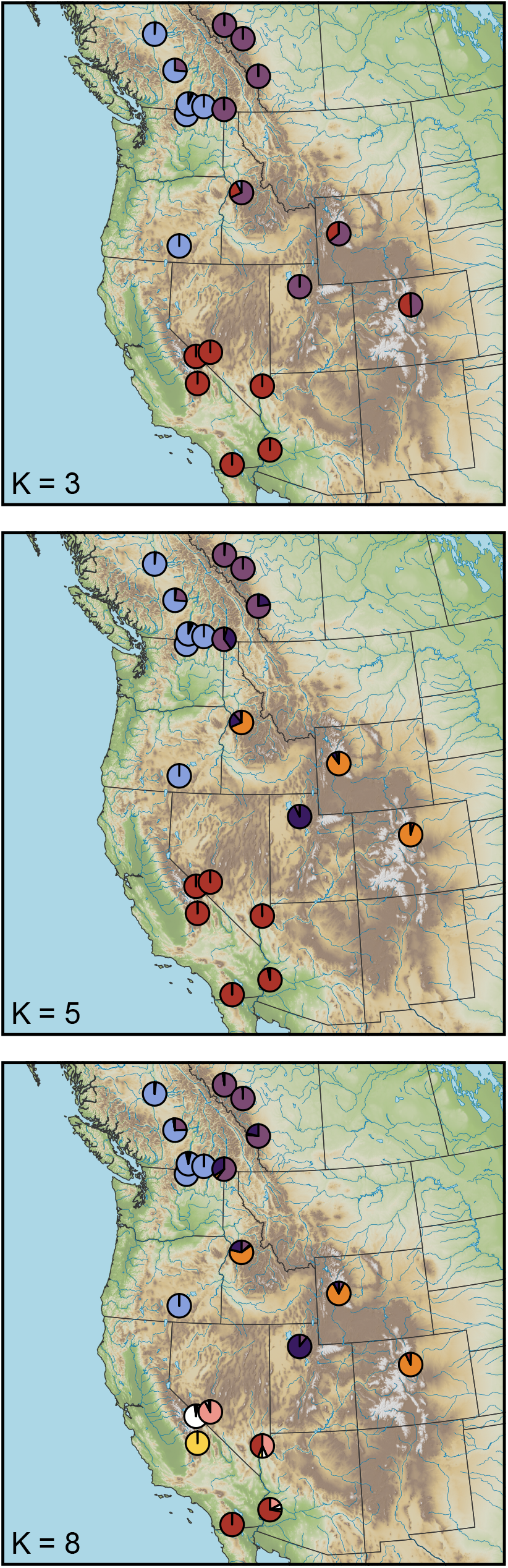
Patterns of hierarchical genetic structure for remnant bighorn sheep populations across western North America. Population mean ancestry coefficients (*q*) are shown for `entropy` models with 3, 5, and 8 genetic clusters (K), each of which are depicted using a different color. Individuals depict previous translocation events, with arrowheads showing the direction of individual movement. See Fig. 2 for barplots displaying individual ancestry for K = 2 – 8.

**Figure 4:**
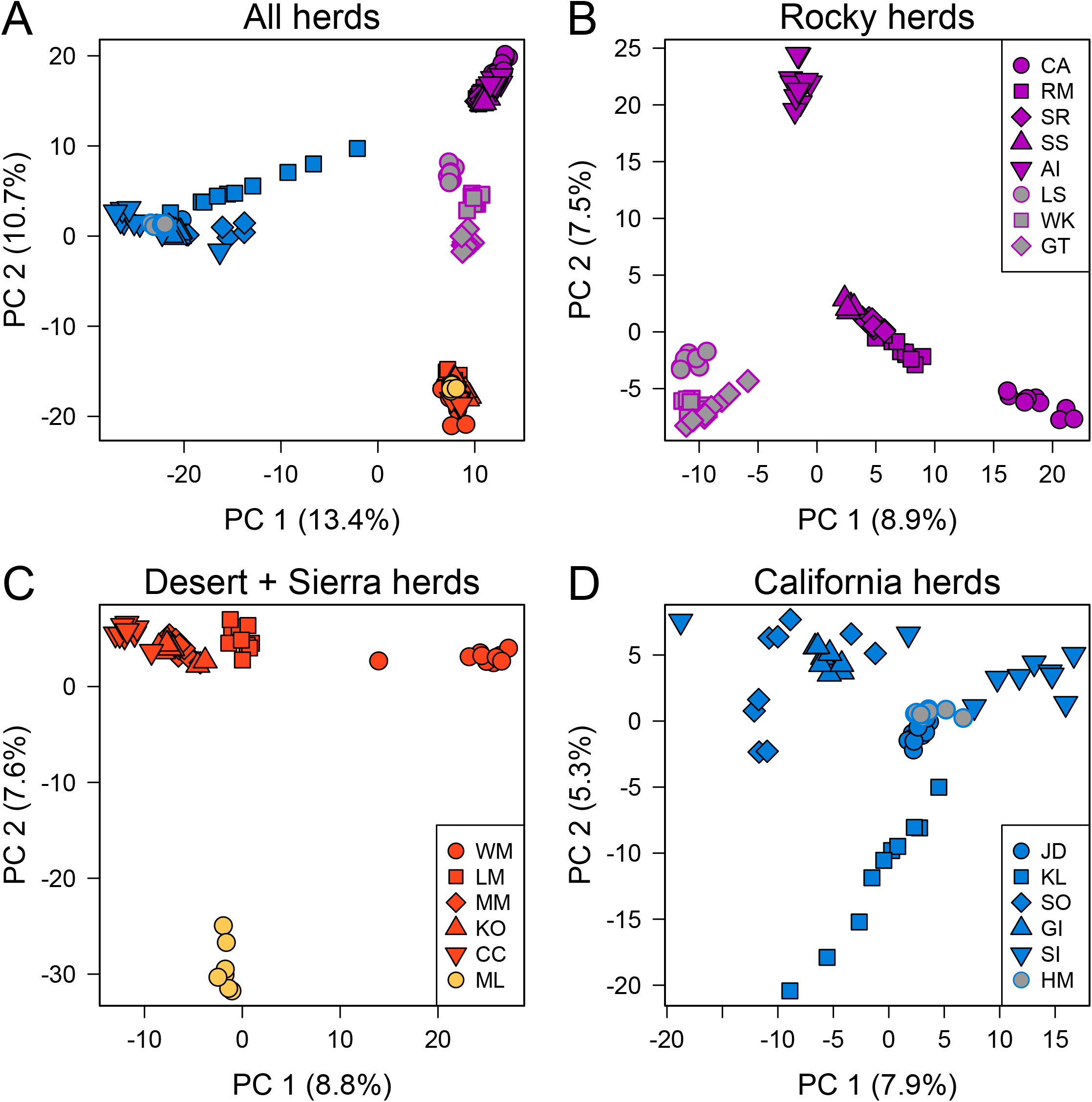
Patterns of population structure among (A) and within (B–D) bighorn sheep lineages was characterized using principal component analysis based on genotype probabilities from `entropy`. In the California lineage, Kamloops (KL) received translocated Rocky Mountain bighorn in 1927 and 1970. Southern Rocky Mountain populations include LS, WK, and GT.

Across all populations, the largest genetic differentiation was found between a desert and a Rocky Mountain population (*F*_ST_ = 0.124; Lone Mountain, LM and Cadomin, CA) (Table S4). In contrast, the smallest differentiation was found between two California populations, Junction Deer Park (JD, also referred to as Williams Lake in previous publications) and Hart Mountain (HM) (*F*_ST_ = 0.008); HM was founded using individuals from JD (Table S1). In general, differentiation among California populations was low (max *F*_ST_ = 0.030; Kamloops Lake, KL versus Sinlahekin, SI) relative to differentiation estimates within the desert (max *F*_ST_ = 0.078; Carrizo Canyon, CC versus White Mountains, WM) and Rocky Mountain (max *F*_ST_ = 0.071; Cadomin, CA versus Lower Salmon, LS) lineages. A neighbor-joining tree based on pairwise *F*_ST_ (Fig. 5) strongly separated populations into three major groupings (California, desert plus Sierra Nevada, and Rocky Mountain), consistent with the ancestry results (Fig. 2).

**Figure 5:**
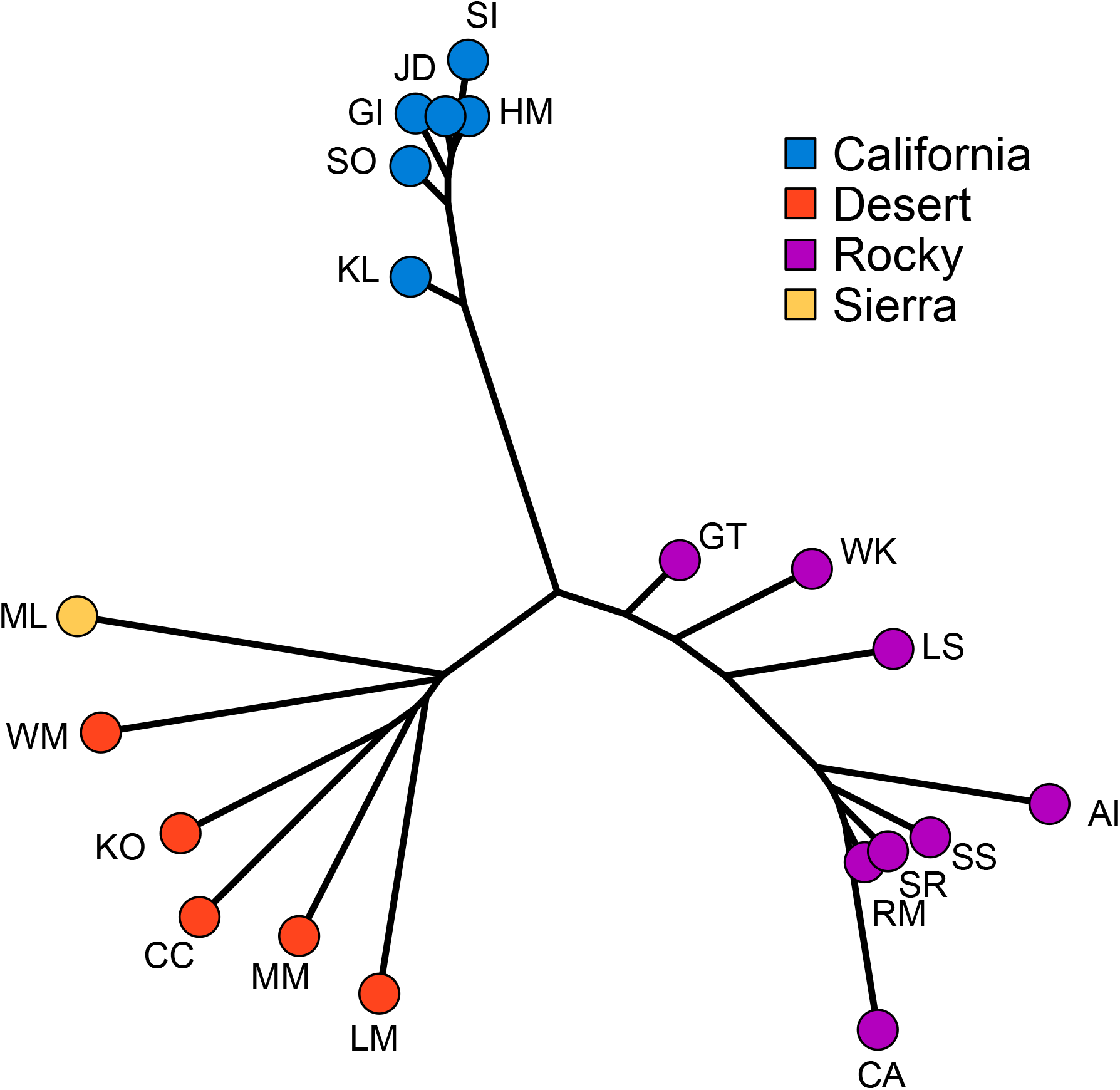
A neighbor-joining tree based on pairwise Hudson’s *F*_ST_ shows patterns of differentiation among populations and lineages. See Table 1 for additional population information.

Levels of genetic diversity, characterized by expected heterozygosity (*H*_E_), ranged from 0.087 (Hart Mountain) to 0.130 (Kofa) (Fig. 6). Estimates were lowest for California populations (*H*_E_ range: 0.087–0.102) and highest for desert populations (*H*_E_ range: 0.112–0.130) and the three more southern Rocky Mountain populations (Georgetown, Lower Salmon, Whiskey Mountain) (*H*_E_ range: 0.126–0.128). The other more northern Rocky Mountain populations (*H*_E_ range: 0.101–0.110) and the sole Sierra Nevada population (*H*_E_ = 0.103) had estimates that were roughly intermediate between California and desert populations.

**Figure 6:**
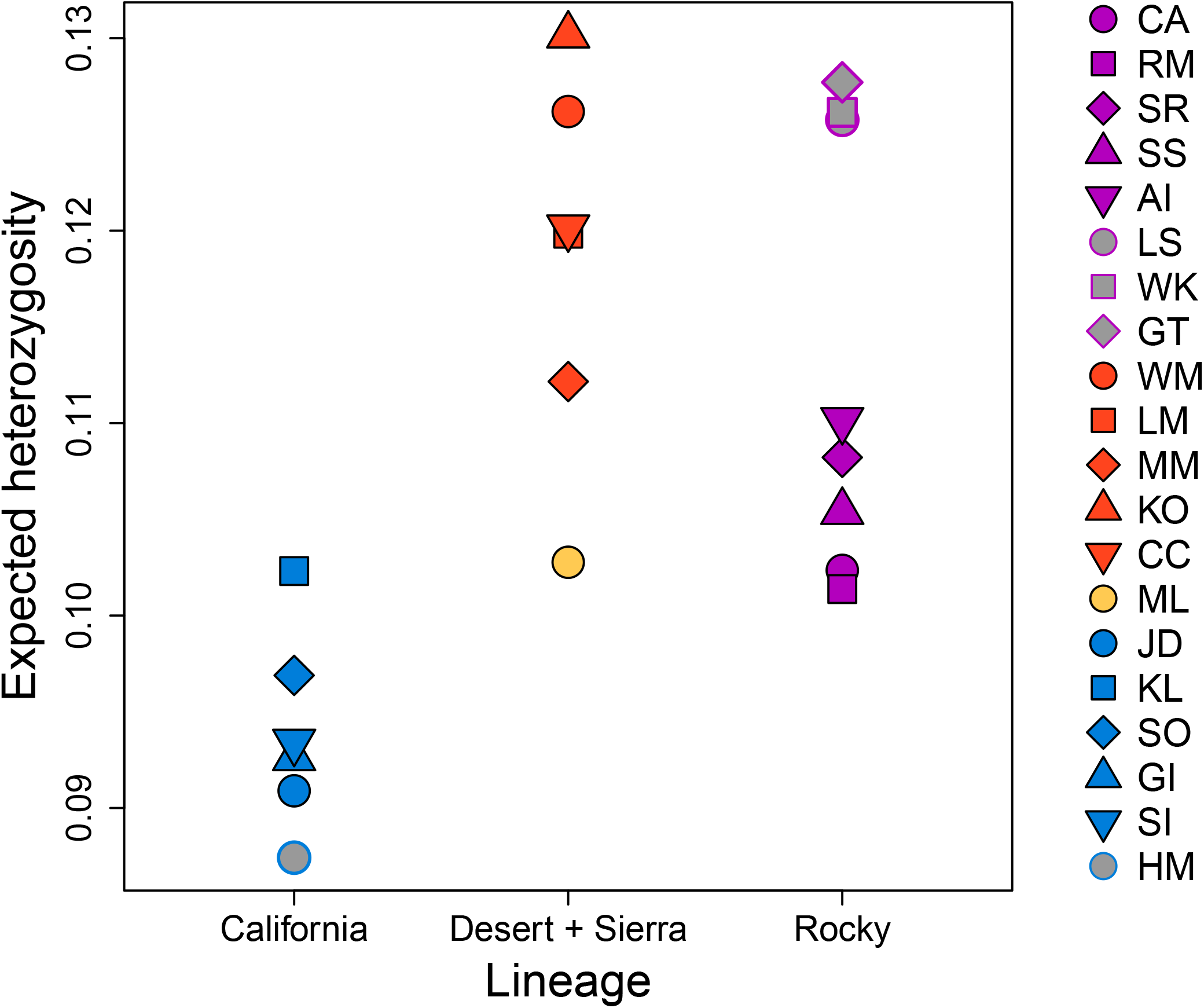
Expected heterozygosity (*H*_E_) estimates for all bighorn sheep populations. Kamloops (KL) has both California and Rocky Mountain ancestry due to translocation.

The maximum likelihood phylogenetic tree had strong bootstrap (bs) support for three major groups: California bighorn sheep, desert plus Sierra Nevada bighorn sheep, and most of the Rocky Mountain populations (Fig. 7). A branch containing most of the Rocky Mountain populations was the earliest split in the tree. However, individuals from more southern Rocky Mountain populations (Georgetown, GT; Lower Salmon, LS; Whiskey, WK) did not have strong support for inclusion in this group. A branch containing GT and LS was placed in between the northern Rocky Mountain branch and the California plus desert plus Sierra Nevada branch with fairly strong support (bs = 88), whereas WK was placed as the earliest diverging population within the the northern Rocky Mountain branch, albeit with low support (bs = 65). California bighorn sheep were sister (i.e., most closely related) to the desert plus Sierra Nevada grouping with strong node support (bs = 97). The California bighorn branch was well supported (bs = 100), although node support within that branch was relatively low and individuals from the same population were not always sister to each other. Within the strongly supported desert plus Sierra Nevada branch (bs = 100), the Sierra Nevada individuals diverged first in a strongly supported branch (bs = 100).

**Figure 7:**
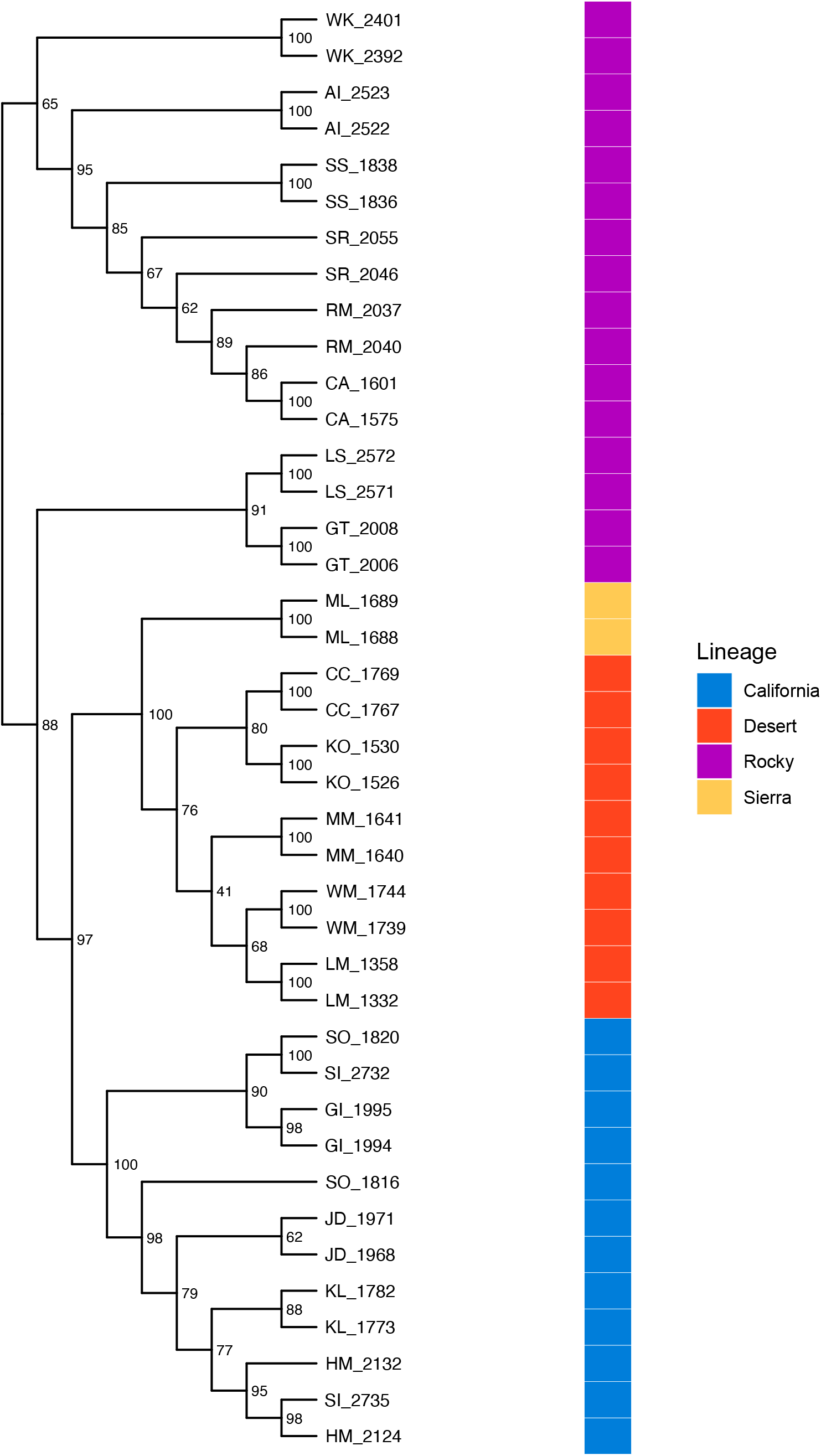
Divergence across lineages and populations is shown with a maximum likelihood phylogenetic tree that was reconstructed using IQ-TREE. Tip labels correspond to population and individual ids, and node labels represent confidence levels based on 1,000 ultrafast bootstraps. See Table 1 for additional population information.

## Discussion

### The history of bighorn sheep diversification

Our results provide strong support for the genetic distinctiveness of the no longer putative California bighorn sheep lineage. In contrast to previous work, including both morphometric and genetic analyses (Ramey, 1993; Wehausen & Ramey, 2000), we determined that populations of bighorn in or sourced from western B. C., previously included in *O. c. californiana* by Cowan (1940), form a well-supported lineage that is at least as deeply differentiated as the Rocky Mountain and desert bighorn subspecies (Fig. 7). We refer hereafter to that lineage, now comprised of bighorn sheep in western B. C. (except where translocated Rocky Mountain bighorn have been established) and translocated populations in the U. S. derived from western B. C., as the California bighorn sheep lineage, although it does not include all populations identified by Cowan (1940) as *O. c. californiana*. Future management of the California lineage would benefit from a more geographically precise nomenclature (e.g., Kokkonen *et al*., 2024), perhaps after further work to clarify the original extent of that lineage.

In our hierarchical analysis of population genetic structure, the California lineage was the first to split from the rest of the putative lineages and populations (Figs. 2, 4). The phylogenetic analysis also exhibited strong support for this split (Fig. 7). Most importantly, our findings clearly supported the hypothesis that the western B. C. portion of the California subspecies are distinct from populations in this study that were described by Cowan (1940) as Rocky Mountain bighorn, refuting the alternate hypothesis of clinal variation across the northern range of the species. Our conclusion is consistent with the findings of Barbosa *et al*. (2021) who analyzed nuclear DNA markers (microsatellites) on a narrower geographic scope and largely from translocated populations, but were in contrast to those of Ramey‘s (1993) analysis of mitochondrial DNA restriction fragment length polymorphisms, wherein the mtDNA haplotype described in the single California bighorn population included in that analysis was also present in northern populations of Rocky Mountain bighorn. That finding may reflect the difference between the history of the nuclear genome and that of the mitochondrial genome (i.e., mitonuclear discordance), an example of mitochondrial capture from some ancient gene flow among lineages, or the lower resolution of RFLP datasets. It is worth noting that even thinhorn and bighorn sheep are paraphyletic for mtDNA (Loehr *et al*., 2006). Morphometric analyses by Wehausen & Ramey (2000) of variation in bighorn skulls had the advantage that skulls from extirpated portions of the range of Cowan’s California subspecies could be included, despite lack of extant native populations as sources for modern DNA. Their study concluded that Rocky Mountain and northern (non-Sierra Nevada) populations of California bighorn did not meet their criteria for subspecies-level distinction, but did note “partial affinity” of California bighorn from British Columbia and Washington with desert bighorn from the Great Basin, and found that skulls from Oregon clearly grouped with the Great Basin desert form. Because we used only extant populations as sources of samples to generate our SNP genotypes, we were not able to obtain data from native bighorn sheep populations present in Oregon, Washington, or northwestern Nevada in the 19th and early 20th centuries that are now extirpated.

Although our results provide clear support for the genetic identity of the California bighorn sheep lineage, less genetic differentiation exists among California bighorn populations compared to Rocky Mountain or desert populations, likely due to the California lineage’s long history of serial translocations from a small pool of donor populations (Table S1; Wild Sheep Working Group, 2015). Signatures of hierarchical population structure did not emerge in the PCA (Fig. 4D) or in `entropy` models with increasing numbers of genetic ancestries (Figs. 2, S1), resulting in relatively small pairwise *F*_ST_ estimates (Fig. 5; Table S4), though at least one split in the phylogenetic tree had strong support (Fig. 7). This suggests that even if managers select the most genetically differentiated California populations as sources for translocations, the potential for genetic augmentation will be limited relative to other lineages. Reduced range-wide differentiation for California bighorn sheep was also recovered by Barbosa *et al*. (2021) based on a microsatellite dataset, though they did report potential evidence for hierarchical structure. The largest estimates of differentiation among California populations in our study stemmed from the presence of Rocky Mountain ancestry in Kamloops Lake (KL; Fig. 2). This Rocky Mountain ancestry is the genetic legacy of translocations of Rocky Mountain bighorn sheep from Banff and Jasper National Parks into the core historic range of California bighorn sheep in 1927 and 1970, respectively (Wild Sheep Working Group, 2015). Rocky Mountain ancestry has now been reported in several populations in the southern Fraser metapopulation (Kamloops Lake in this study, Fig. 2; Fraser-River Churn Creek and Spences Bridge in Barbosa *et al*., 2021), but fortunately not in the Junction Deer Park (JD) population that served as the primary translocation source for most of the reestablished California bighorn populations in the United States (Table S1; Wild Sheep Working Group, 2015; Barbosa *et al*., 2021). Continued spread of Rocky Mountain ancestry throughout this metapopulation will limit the options for future California bighorn translocations, compounding the challenges of managing this lineage. The main take home point is that the history of translocations in both Canada and the United States has greatly complicated the situation, and that managers are now left in a place with few good choices. We believe this provides a fair and balanced summary of the situation on the ground for California bighorn sheep, and hope this will stimulate future conversations, management actions, and genetic studies to further explore options for this lineage.

Divergence within Rocky Mountain bighorn sheep largely coincides with a history of isolation-by-distance along the north to south axis of the Rocky Mountains. This pattern is readily apparent in the `entropy` ancestry assignments for models with K ≥ 5, where ancestry shifts from purple to dark blue to orange in a clinal pattern across descending latitude (Figs. 2 & 3; note that the Antelope Island population was founded with individuals from Montana; Table S1). Isolation-by-distance has been reported in previous assessments of Rocky Mountain bighorn sheep genetic structure, particularly for Canadian populations that have been shielded from translocations more than their U. S. counterparts (Deakin *et al*., 2020; Flesch *et al*., 2020; Love Stowell *et al*., 2020). Given the geographic gaps in our sampling scheme, we are unable to discern the relative influence that prominent barriers to gene flow (e.g., large rivers, low elevation valleys) have had in shaping population structure, but these factors have been implicated in other studies (Deakin *et al*., 2020). Despite this, our results are consistent with a major geographic split between more northern and southern Rocky Mountain populations, as evidenced by clustering in the PCA (Fig. 4A,B), large pairwise *F*_ST_ estimates (Fig. 5, Table S4), and the supported branch on the phylogenetic tree containing Lower Salmon and Georgetown individuals (LS and GT; Fig. 7). Due to relative proximity of southern Rocky Mountain populations to areas likely occupied historically by desert bighorn, this may also reflect historical gene flow among those lineages. More fine-scale spatial sampling is needed to further clarify this possibility, though the dearth of remnant populations in the U. S. may limit potential insights.

Our sampling of desert bighorn sheep included nearly all currently or previously recognized southern lineages (based on Buchalski *et al*., 2016), with Lone Mountain (LM), the Muddy Mountains (MM), and the White Mountains (WM) representing Nelson’s desert bighorn (the sole currently recognized desert subspecies), Kofa (KO) representing Mexican desert bighorn (*O. c. mexicana*, a previously recognized subspecies), and Carrizo Canyon (CC) representing peninsular desert bighorn (*O. c. cremnobates*, a previously recognized subspecies that is now managed as a federally endangered distinct population segment; U. S. Fish and Wildlife Service, 1998). The degree of differentiation among desert bighorn populations was large relative to the other lineages (*F*_ST_ = 0.057 – 0.078; Fig. 5), consistent with the results of Buchalski *et al*. (2016) based on microsatellites and mtDNA. While Buchalski *et al*. (2016) found that Nelson, Mexican, and Peninsular populations clearly split into distinct ancestry clusters, the K = 8 `entropy` model in this study recovered a north to south step-wise transition of ancestry across those populations (Figs. 2 & 3). Previous studies have raised the possibility that Lone Mountain (and herds that have received translocations from Lone Mountain) might represent a relict Great Basin lineage of desert bighorn sheep based on genetic (Jahner *et al*., 2019; Wright *et al*., 2024b) and morphological (Wehausen, 1991) differences from Mojave Desert populations (e.g., the Muddy Mountains). The genetic results in this study were ambiguous about the relationship of Lone Mountain with other populations: the phylogenetic analysis placed Lone Mountain sister to the White Mountains (albeit with modest support; Fig. 7), whereas the population genetic analyses allied Lone Mountain with the other more southern desert populations (Figs. 2, 5). As such, a final verdict on the possibility of a Great Basin lineage of desert bighorn sheep will await a future study with expanded genomic sampling.

Our analysis confirmed that Sierra Nevada bighorn formed a lineage that was strongly distinct from desert bighorn, yet one that was more closely related to desert bighorn than relationships among desert, Rocky Mountain, and California lineages (Fig. 7). Sierra Nevada bighorn are morphologically distinct from desert bighorn sheep (Wehausen & Ramey, 2000), likely due to adaptations to extreme cold in the high elevations of the Sierra Nevada (*>*4,000m) versus extreme heat experienced by many populations of desert bighorn. Ramey (1993) conducted the most geographically complete published evaluation of genetic variation (mitochondrial RFLPs) in bighorn sheep previous to this study and concluded that Sierra Nevada bighorn were distinct from, yet more closely related to, desert bighorn than to other lineages. In contrast, phylogenies based on mitochondrial sequence data were more ambiguous as to the placement of Sierra Nevada bighorn relative to other lineages, likely due to discordance between mitochondrial and nuclear genomes, small sample sizes, and the limitations of relatively short sections of control region sequence in making inference over deep time scales (note that only one Sierra Nevada individual was included in those analyses; Buchalski *et al*., 2016; Wright *et al*., 2024a). Buchalski et al. (2016) also used discriminant analysis of principal components on microsatellite genotypes and found that Sierra Nevada bighorn were as strongly differentiated as Rocky Mountain and desert bighorn. Given the rapid evolution of microsatellite markers within a typically constrained range of allele sizes, that result may represent more recent time frames than our dataset. Sierra Nevada bighorn are geographically close to desert bighorn sheep, and occasional movements of individuals between the two subspecies’ ranges were reported historically (Jones, 1950) and in one recent instance (Epps *et al*., 2024). Desert bighorn in the White Mountains, which separated from other desert bighorn in our analysis in the first hierarchical subdivision (Fig. 2), exhibit seasonal altitudinal migration to high elevation habitat (*>*4,000m) and are exposed to cold temperatures (Wehausen, 1983). Wehausen & Ramey (1993) distinguished the southern (hot desert) desert bighorn from the northern (cold desert, including White Mountains, Death Valley, and extirpated populations to the north) desert bighorn on the basis of skull morphology. Thus, the phylogeographic history of Sierra Nevada and desert bighorn sheep in the northern portion of the range, including many now-extirpated populations in the Great Basin, is likely complex and deserving of finer-scaled investigation.

### Informing future management

Our findings that the western British Columbia portion of Cowan‘s (1940) California subspecies is as deeply differentiated as the currently recognized desert and Rocky Mountain subspecies, and more differentiated than the relationship between desert and Sierra Nevada subspecies, has important implications for management. Translocated populations derived from western B. C. are widespread across Washington, Oregon, southwestern Idaho, northwestern Nevada, and present in Utah and North Dakota. Most U.S. states, and B. C. at this time, still manage populations of Rocky Mountain versus western B. C. lineage separately, and in most cases, have avoided mixing them. Studies of a mixed population established in North Dakota found individuals with the more regionally-appropriate Rocky Mountain bighorn ancestry had higher fitness than those of western B. C. (California) ancestry (Wiedmann & Sargeant, 2014; Bleich *et al*., 2018), further highlighting the need to correctly identify divergent lineages below the species level that may exhibit different adaptations. We recommend that managers avoid mixing those lineages in the future, and note that the translocated populations of western B. C. ancestry now present in Washington, Oregon, northwestern Nevada, southwestern Idaho, and several locations in Utah may represent an important portion of the remaining gene pool for this lineage, even though the low genetic diversity of many of these populations arising from multiple founder effects is well known (Whittaker *et al*., 2004; Olson *et al*., 2013; Barbosa *et al*., 2021; Spaan *et al*., 2021). We suggest that detailed investigation of those populations might determine whether new strategies could be developed for preserving that remaining genetic diversity. We also reiterate the need to determine the lineage of extirpated populations in northern Nevada, Oregon, and Washington. Wehausen & Ramey (2000) noted that skulls exhibited desert bighorn characteristics as far north as the Columbia River, and no clear barriers to gene flow from desert bighorn native to the south are apparent, suggesting that California bighorn now present in the Great Basin deserts because of translocations may not be best adapted to those arid habitats (Malaney *et al*., 2015).

DNA sequencing data has become widely available for conservation and management guidance (Hohenlohe *et al*., 2021; Allendorf *et al*., 2022), allowing key insights into previously understudied taxa. This wealth of information will be particularly useful for managers considering translocations to augment existing populations by allowing for better identification of potential recipient populations, as well as the selection of the best donor populations and individuals. In some cases, managers may be best served by selecting individuals that are expected to contribute new genetic variation to the recipient population (recognizing that genetic variation is just one of many important variables to consider). However, an open question in conservation genetics is whether the potential benefits gained by translocations involving well-differentiated populations (i.e., different evolutionary significant units) outweigh the potential costs of disrupting local adaptation via outcrossing (Weeks *et al*., 2011; Bell *et al*., 2019; Liddell *et al*., 2021). While we cannot offer a resolution to this decision for bighorn sheep, to protect the remaining natural variation that still exists on the landscape, we suggest that managers should avoid translocating individuals from different genetic lineages into or to within dispersal distance of remnant, native populations whenever possible. Likewise, continued monitoring efforts should be designed to provide information on the fitness consequences of translocations relative to subsequent diet, physiology, survival, and reproductive success of translocated individuals in order to augment the effectiveness of future efforts. For bighorn sheep and many other species, translocation and assisted migration will become an increasingly important conservation management tool as these systems face rapid environmental change.

## Acknowledgments

We thank: Alex Buerkle for valuable feedback on several aspects of the project; Katie Wagner for critical advice on the phylogenetic analyses; Seth Romero for guidance on creating beautiful maps; three anonymous reviewers, Sean Harrington, Sam Johnson, Joel Ogwang, Maria Paula Rodriguez, Will Rosenthal, and Drew Suchomel for providing comments on an earlier draft. Samples and logistical support were generously provided by Hank Edwards, Marco Festa-Bianchet, Nathan Galloway, Bill Jex, Doug McWhirter, Riley Peck, Rusty Robinson, Eric Rominger, Caeley Thacker, Don Whittaker, Brett Wiedmann, Shari Willmott, Mary Wood, Alberta Fish and Wildlife, British Columbia Ministry of Water, Land, and Resource Stewardship, California Department of Fish and Wildlife, Colorado Parks and Wildlife, Idaho Department of Fish and Game, Nevada Department of Wildlife, New Mexico Department of Game and Fish, North Dakota Game and Fish Department, Oregon Department of Fish and Wildlife, Utah Division of Wildlife Resources, Washington Department of Fish and Wildlife, and Wyoming Game and Fish Department. This program is supported by Federal financial assistance titled “Statewide Game Management” submitted to the U.S. Fish and Wildlife Service’s CFDA Program 15.611 and is made under the authority of: Pittman-Robertson Wildlife Restoration Act of 1937, 16 U.S.C. 669-669k. This research was supported by funding from the Nevada Department of Wildlife, Idaho Department of Fish and Game, and the Wild Sheep Foundation. JPJ was supported by the Modelscape Consortium with funding from NSF (OIA-2019528); MDM and TLP were supported in part by NSF (OIA-1826801). Analyses were conducted on the University of Wyoming’s Advanced Research Computing Center and its Beartooth Computing Environment, Intel x86_64 cluster (https://doi.org/10.15786/M2FY47). Any opinions, findings, conclusions, or recommendations expressed in this publication are those of the authors and do not necessarily reflect the official views of state and federal agencies, including the Nevada Department of Wildlife, the U.S. Fish and Wildlife Service, or the U.S. Department of Agriculture.

## Data Archiving Statement

The raw data associated with this manuscript will be submitted to the Dryad Digital Repository after manuscript is accepted for publication.

## References

Allendorf FW, Funk WC, Aitken SN, Byrne M, Luikart G (2022) Conservation and the genomics of populations. 3rd edn., Oxford University Press.

Barbosa S, Andrews KR, Harris RB, et al. (2021) Genetic diversity and divergence among bighorn sheep from reintroduced herds in Washington and Idaho. The Journal of Wildlife Management, 85, 1214–1231.

Bell DA, Robinson ZL, Funk WC, et al. (2019) The exciting potential and remaining uncertainties of genetic rescue. Trends in Ecology & Evolution, 34, 1070–1079.

Bleich V, Wehausen J, Torres S, Anderson K, Stephenson T (2021) Fifty years of bighorn sheep translocations: details from California (1971–2020). Desert Bighorn Council Transactions, 56, 1–32.

Bleich VC, Sargeant GA, Wiedmann BP (2018) Ecotypic variation in population dynamics of reintroduced bighorn sheep. The Journal of Wildlife Management, 82, 8–18.

Buchalski MR, Navarro AY, Boyce WM, et al. (2015) Genetic population structure of Peninsular bighorn sheep (Ovis canadensis nelsoni) indicates substantial gene flow across US–Mexico border. Biological Conservation, 184, 218–228.

Buchalski MR, Sacks BN, Gille DA, et al. (2016) Phylogeographic and population genetic structure of bighorn sheep (Ovis canadensis) in North American deserts. Journal of Mammalogy, 97, 823–838.

Buechner HK (1960) The bighorn sheep in the United States, its past, present, and future. Wildlife Monographs, 4, 1–174.

Cassirer EF, Manlove KR, Almberg ES, et al. (2018) Pneumonia in bighorn sheep: Risk and resilience. The Journal of Wildlife Management, 82, 32–45.

Chafin TK, Zbinden ZD, Douglas MR, et al. (2021) Spatial population genetics in heavily managed species: Separating patterns of historical translocation from contemporary gene flow in white-tailed deer. Evolutionary Applications, 14, 1673–1689.

Clavero M, Naves J, Lucena-Perez M, Revilla E (2024) Taxonomic inflation as a conservation trap for inbred populations. Evolutionary Applications, 17, e13677.

Cowan IM (1940) Distribution and variation in the native sheep of North America. The American Midland Naturalist, 24, 505–580.

Creech TG, Epps CW, Wehausen JD, et al. (2020) Genetic and environmental indicators of climate change vulnerability for desert bighorn sheep. Frontiers in Ecology and Evolution, 8, 279.

Danecek P, Auton A, Abecasis G, et al. (2011) The variant call format and vcftools. Bioinformatics, 27, 2156–2158.

Deakin S, Gorrell JC, Kneteman J, Hik DS, Jobin R, Coltman D (2020) Spatial genetic structure of Rocky Mountain bighorn sheep Ovis canadensis canadensis at the northern limit of their native range. Canadian Journal of Zoology, 98, 317–330.

Epps CW, Buchalski M, Jahner JP, Sim Z (2026) Application of genetics to taxonomy, biology, conservation, and management. In: Mountain sheep in North America: biology, ecology, conservation, and management (eds. Krausman PR, Jex B), CRC Press.

Epps CW, Buchalski MR, Crowhurst R, Aiello CM (2024) Evaluation of foraying ram genetics relative to neighboring desert bighorn populations. Report to California Department of Fish and Wildlife.

Epps CW, Crowhurst RS, Nickerson BS (2018) Assessing changes in functional connectivity in a desert bighorn sheep metapopulation after two generations. Molecular Ecology, 27, 2334–2346.

Epps CW, Palsbøll PJ, Wehausen JD, Roderick GK, Ramey RR, McCullough DR (2005) Highways block gene flow and cause a rapid decline in genetic diversity of desert bighorn sheep. Ecology Letters, 8, 1029–1038.

Epps CW, Wehausen JD, Palsbøll PJ, McCullough DR (2010) Using genetic tools to track desert bighorn sheep colonizations. The Journal of Wildlife Management, 74, 522–531.

Flesch EP, Graves TA, Thomson JM, et al. (2020) Evaluating wildlife translocations using genomics: A bighorn sheep case study. Ecology and Evolution, 10, 13687–13704.

Fraser DJ, Bernatchez L (2001) Adaptive evolutionary conservation: towards a unified concept for defining conservation units. Molecular Ecology, 10, 2741–2752.

Gann WJ, Gray SS, Dittmar RO, Gonzalez CE, Harveson LA (2020) Effective pronghorn translocation methodology: a long-term summary. Wildlife Society Bulletin, 44, 599–609.

Garcia de Leaniz C, Fleming I, Einum S, et al. (2007) A critical review of adaptive genetic variation in Atlantic salmon: implications for conservation. Biological Reviews, 82, 173–211.

Geist V (1971) Mountain sheep: a study in behavior and evolution. University of Chicago Press, Chicago, IL.

Geist V (1985) On Pleistocene bighorn sheep: some problems of adaptation, and relevance to today’s american megafauna. Wildlife Society Bulletin, 13, 351–359.

Gelman A, Rubin DB (1992) Inference from iterative simulation using multiple sequences. Statistical Science, 7, 457–472.

Gille DA, Buchalski MR, Conrad D, et al. (2019) Genetic outcomes of translocation of bighorn sheep in Arizona. The Journal of Wildlife Management, 83, 838–854.

Gompert Z, Lucas LK, Buerkle CA, Forister ML, Fordyce JA, Nice CC (2014) Admixture and the organization of genetic diversity in a butterfly species complex revealed through common and rare genetic variants. Molecular Ecology, 23, 4555–4573.

Grinnell J (1912) The bighorn of the Sierra Nevada. University of California Publications in Zoology, 10, 143–153.

Hendry AP, Lohmann LG, Conti E, et al. (2010) Evolutionary biology in biodiversity science, conservation, and policy: a call to action. Evolution, 64, 1517–1528.

Hoang DT, Chernomor O, Von Haeseler A, Minh BQ, Vinh LS (2018) UFBoot2: improving the ultrafast bootstrap approximation. Molecular Biology and Evolution, 35, 518–522.

Hohenlohe PA, Funk WC, Rajora OP (2021) Population genomics for wildlife conservation and management. Molecular Ecology, 30, 62–82.

Hudson RR, Slatkin M, Maddison WP (1992) Estimation of levels of gene flow from DNA sequence data. Genetics, 132, 583–589.

IUCN/SSC (2013) Guidelines for reintroductions and other conservation translocations. Version 1.0. IUCN Species Survival Commission, Gland, Switzerland.

Jahner JP, Matocq MD, Malaney JL, et al. (2019) The genetic legacy of 50 years of desert bighorn sheep translocations. Evolutionary Applications, 12, 198–213.

[dataset] Jahner JP, Parchman TL, Matocq MD, et al. (2025) Data from: Resolving the evolutionary history of bighorn sheep to inform future management. Dryad Digital Repository (doi:XXX).

Jones FL (1950) A survey of the Sierra Nevada bighorn. Sierra Club Bulletin, 35, 29–76.

Kallman H (1987) Restoring America’s Wildlife, 1937-1987: The First 50 Years of the Federal Aid in Wildlife Restoration (Pittman-Robertson) Act. US Department of the Interior, Fish and Wildlife Service.

Kalyaanamoorthy S, Minh BQ, Wong TK, Von Haeseler A, Jermiin LS (2017) ModelFinder: fast model selection for accurate phylogenetic estimates. Nature Methods, 14, 587–589.

Kokkonen AL, Searle PC, Shiozawa DK, Evans RP (2024) Using de novo transcriptomes to decipher the relationships in cutthroat trout subspecies (Oncorhynchus clarkii). Evolutionary Applications, 17, e13735.

Langmead B, Salzberg SL (2012) Fast gapped-read alignment with Bowtie 2. Nature Methods, 9, 357.

Li H, Durbin R (2009) Fast and accurate short read alignment with Burrows–Wheeler transform. Bioinformatics, 25, 1754–1760.

Li H, Handsaker B, Wysoker A, et al. (2009) The sequence alignment/map format and SAMtools. Bioinformatics, 25, 2078–2079.

Liddell E, Sunnucks P, Cook CN (2021) To mix or not to mix gene pools for threatened species management? few studies use genetic data to examine the risks of both actions, but failing to do so leads disproportionately to recommendations for separate management. Biological Conservation, 256, 109072.

Loehr J, Worley K, Grapputo A, Carey J, Veitch A, Coltman D (2006) Evidence for cryptic glacial refugia from North American mountain sheep mitochondrial DNA. Journal of Evolutionary Biology, 19, 419–430.

Love Stowell SM, Gagne RB, McWhirter D, Edwards W, Ernest HB (2020) Bighorn sheep genetic structure in Wyoming reflects geography and management. The Journal of Wildlife Management, 84, 1072–1090.

Luikart G, Allendorf FW (1996) Mitochondrial-DNA variation and genetic-population structure in Rocky Mountain bighorn sheep (Ovis canadensis canadensis). Journal of Mammalogy, 77, 109–123.

Malaney JL, Feldman CR, Cox M, Wolff P, Wehausen JD, Matocq MD (2015) Translocated to the fringe: genetic and niche variation in bighorn sheep of the Great Basin and northern Mojave deserts. Diversity and Distributions, 21, 1063–1074.

Martin LD, Gilbert BM (1978) Excavations at Natural Trap Cave. Transactions of the Nebraska Academy of Sciences and Affiliated Societies, 336.

Miller J, Poissant J, Hogg J, Coltman D (2012) Genomic consequences of genetic rescue in an insular population of bighorn sheep (Ovis canadensis). Molecular Ecology, 21, 1583–1596.

Miller JM, Moore SS, Stothard P, Liao X, Coltman DW (2015) Harnessing cross-species alignment to discover SNPs and generate a draft genome sequence of a bighorn sheep (Ovis canadensis). BMC Genomics, 16, 1–8.

Minh BQ, Nguyen MAT, Von Haeseler A (2013) Ultrafast approximation for phylogenetic bootstrap. Molecular Biology and Evolution, 30, 1188–1195.

Nguyen LT, Schmidt HA, Von Haeseler A, Minh BQ (2015) IQ-TREE: a fast and effective stochastic algorithm for estimating maximum-likelihood phylogenies. Molecular Biology and Evolution, 32, 268–274.

Olson ZH, Whittaker DG, Rhodes Jr OE (2012) Evaluation of experimental genetic management in reintroduced bighorn sheep. Ecology and Evolution, 2, 429–443.

Olson ZH, Whittaker DG, Rhodes Jr OE (2013) Translocation history and genetic diversity in reintroduced bighorn sheep. The Journal of Wildlife Management, 77, 1553–1563.

Ord TJ, Summers TC (2015) Repeated evolution and the impact of evolutionary history on adaptation. BMC Evolutionary Biology, 15, 1–12.

Paradis E, Schliep K (2019) ape 5.0: an environment for modern phylogenetics and evolutionary analyses in R. Bioinformatics, 35, 526–528.

Parchman TL, Gompert Z, Mudge J, Schilkey FD, Benkman CW, Buerkle CA (2012) Genome-wide association genetics of an adaptive trait in lodgepole pine. Molecular Ecology, 21, 2991–3005.

R Core Team (2022) R: a language and environment for statistical computing.. R Foundation for Statistical Computing, Vienna, Austria. URL https://www.R-project.org/.

Ramey RR (1993) Evolutionary genetics and systematics of North American mountain sheep: implications for conservation. Ph.D. thesis, Cornell University.

Ramey RR (1995) Mitochondrial DNA variation, population structure, and evolution of mountain sheep in the south-western United States and Mexico. Molecular Ecology, 4, 429–440.

Sacks BN, Davis TM, Batter TJ (2024) Genetic structure of California’s elk: a legacy of extirpations, reintroductions, population expansions, and admixture. The Journal of Wildlife Management, 88, e22539.

Santos SH, Peery RM, Miller JM, et al. (2021) Ancient hybridization patterns between bighorn and thinhorn sheep. Molecular Ecology, 30, 6273–6288.

Shastry V, Adams PE, Lindtke D, et al. (2021) Model-based genotype and ancestry estimation for potential hybrids with mixed-ploidy. Molecular Ecology Resources, 21, 1434–1451.

Singer FJ, Papouchis CM, Symonds KK (2000) Translocations as a tool for restoring populations of bighorn sheep. Restoration Ecology, 8, 6–13.

Spaan RS, Epps CW, Crowhurst R, Whittaker D, Cox M, Duarte A (2021) Impact of Mycoplasma ovipneumoniae on juvenile bighorn sheep (Ovis canadensis) survival in the northern Basin and Range ecosystem. PeerJ, 9, e10710.

Turbek SP, Funk WC, Ruegg KC (2023) Where to draw the line? expanding the delineation of conservation units to highly mobile taxa. Journal of Heredity, 114, 300–311.

U. S. Fish and Wildlife Service (1998) Endangered and threatened wildlife and plants; endangered status for the peninsular ranges population segment of the desert bighorn sheep in Southern California. Federal Register, 63, 13134–13150.

Wagner CE, Keller I, Wittwer S, et al. (2013) Genome-wide RAD sequence data provide unprecedented resolution of species boundaries and relationships in the Lake Victoria cichlid adaptive radiation. Molecular Ecology, 22, 787–798.

Wang X (1988) Systematics and population ecology of late Pleistocene bighorn sheep (Ovis canadensis) of Natural Trap Cave, Wyoming. Transactions of the Nebraska Academy of Sciences and Affiliated Societies, 194.

Weeks AR, Sgro CM, Young AG, et al. (2011) Assessing the benefits and risks of translocations in changing environments: a genetic perspective. Evolutionary Applications, 4, 709–725.

Wehausen JD (1983) White Mountain bighorn sheep: an analysis of current knowledge and management alternatives. Inyo National Forest Administrative Report.

Wehausen JD (1991) Some potentially adaptive characters of mountain sheep populations in the Owens Valley region. In: Natural history of eastern California and high-altitude research (eds. Hall C, Doyle-Jones V, Widawski B), pp. 256–267, White Mountain Research Station, Los Angeles, CA.

Wehausen JD, Bleich VC, Ramey RR, et al. (2005) Correct nomenclature for Sierra Nevada bighorn sheep. California Fish and Game, 91, 216.

Wehausen JD, Kelley ST, Ramey RR (2011) Domestic sheep, bighorn sheep, and respiratory disease: a review of the experimental evidence. California Fish and Game, 97, 7–24.

Wehausen JD, Ramey RR (1993) A morphometric reevaluation of the Peninsular bighorn subspecies. Desert Bighorn Council Transactions, 37, 1–10.

Wehausen JD, Ramey RR (2000) Cranial morphometric and evolutionary relationships in the northern range of Ovis canadensis. Journal of Mammalogy, 81, 145–161.

Whiting JC, Bleich VC, Bowyer RT, Epps CW (2023) Restoration of bighorn sheep: History, successes, and remaining conservation issues. Frontiers in Ecology and Evolution, 11, 1083350.

Whittaker DG, Ostermann SD, Boyce WM (2004) Genetic variability of reintroduced California bighorn sheep in Oregon. The Journal of Wildlife Management, 68, 850–859.

Wiedmann BP, Sargeant GA (2014) Ecotypic variation in recruitment of reintroduced bighorn sheep: Implications for translocation. The Journal of Wildlife Management, 78, 394–401.

Wild Sheep Working Group (2015) Records of wild sheep translocations-United States and Canada, 1922-present. Western Association of Fish and Wildlife Agencies, USA.

Winter S, Fennessy J, Janke A (2018) Limited introgression supports division of giraffe into four species. Ecology and Evolution, 8, 10156–10165.

Wright EA, Buchalski MR, Bradley RD (2024a) Mitochondrial DNA indicates that extirpated Ovis canadensis texianus was a member of the desert bighorn sheep complex. Occasional Papers of the Museum of Texas Tech University, 390, 1–29.

Wright EA, Manthey JD, Buchalski MR, et al. (2024b) Genomic affinity following restoration of a locally extirpated species: a case study of desert bighorn sheep in Texas. Conservation Genetics, 25, 1209–1230.

Yu G, Smith DK, Zhu H, Guan Y, Lam TTY (2017) ggtree: an R package for visualization and annotation of phylogenetic trees with their covariates and other associated data. Methods in Ecology and Evolution, 8, 28–36.

