## Supplemental figures and tables for "Resolving the evolutionary history of bighorn sheep to inform future management: an answer to the California bighorn lineage question"

| Lineage | Recipient population | Source population | Source lineage | Year | $N$ |
|---|---|---|---|---|---|
| California | Hart Mountain, OR | Junction, BC | California | 1954 | 20 |
| | Kamloops Lake, BC | Junction, BC | California | 1966 | 11 |
| | Sinlahekin, WA | Junction, BC | California | 1957 | 18 |
| | Sinlahekin, WA | Aldrich Mountains, OR | California | 1990 | 4 |
| | Sinlahekin, WA | South Thompson, BC | California | 1996 | 4 |
| | Sinlahekin, WA | Lower John Day, OR | California | 2003 | 8 |
| | South Okanagan, BC | Junction, BC | California | 1952 | 4 |
| Rocky | Antelope Island, UT | Rocky Boys, MT | Rocky | 2020 | 25 |
| | Georgetown, CO | Tarryall Range, CO | Rocky | 1946 | 33 |
| | Georgetown, CO | Tarryall Range, CO | Rocky | 1949 | 14 |
| | Ram Mountain, AB | Cadomin, AB | Rocky | 2004 | 6 |
| | Ram Mountain, AB | Cadomin, AB | Rocky | 2005 | 6 |
| | Ram Mountain, AB | Cadomin, AB | Rocky | 2007 | 12 |
| Sierra | Mount Langley, CA | Sand Mountain, CA | Sierra | 1980 | 4 |
| | Mount Langley, CA | Sawmill Canyon, CA | Sierra | 1980 | 7 |
| | Mount Langley, CA | Sand Mountain, CA | Sierra | 1982 | 6 |
| | Mount Langley, CA | Sawmill Canyon, CA | Sierra | 1982 | 9 |
| | Mount Langley, CA | Sand Mountain, CA | Sierra | 1987 | 2 |

Table S1: Known translocations of bighorn sheep into the populations included in this study, based on the records from Wild Sheep Working Group (2015). The translocations involving Sierra Nevada bighorn sheep have occurred within their small, recently known historical distribution. Introductions of Rocky Mountain bighorn sheep into Hart Mountain in 1939 and California bighorn sheep into Antelope Island in 1997 and 2000 were ultimately unsuccessful and not shown here.

| K | Mean DIC | S.D. DIC |
|---|---|---|
| 1 | 9,041,589 | 94,766 |
| 2 | 8,094,440 | 46,893 |
| 3 | 7,510,411 | 35,900 |
| 4 | 8,365,119 | 242,647 |
| 5 | 9,350,162 | 635,912 |
| 6 | 9,422,079 | 572,386 |
| 7 | 9,288,110 | 235,603 |
| 8 | 9,619,230 | 380,355 |
| 9 | 9,498,826 | 399,098 |
| 10 | 11,971,802 | 2,390,384 |

Table S2: Five replicate chains were run for full dataset `entropy` models including 1 – 10 ancestries (K). Models were evaluated using deviance information criteria (DIC) scores.

| | California | | Desert & Sierra | | Rocky | |
|---|---|---|---|---|---|---|
| K | Mean DIC | S.D. DIC | Mean DIC | S.D. DIC | Mean DIC | S.D. DIC |
| 1 | 2,499,339 | 19,887 | 1,547,912 | 11,130 | 3,519,925 | 26,033 |
| 2 | 3,907,585 | 67,405 | 1,687,776 | 43,446 | 3,421,640 | 45,826 |
| 3 | 3,684,571 | 179,180 | 1,717,751 | 55,250 | 3,493,869 | 58,214 |
| 4 | 5,651,491 | 1,076,900 | 1,761,733 | 31,547 | 3,336,626 | 26,541 |
| 5 | 5,727,455 | 215,545 | 1,524,669 | 11,516 | 3,522,155 | 62,456 |

Table S3: For each data subset (California, desert plus Sierra, Rocky), replicate chains were run for `entropy` models including 1 – 5 ancestries (K). Models were evaluated using deviance information criteria (DIC) scores, which can be compared across Ks but not across different subsets.

| | CA | RM | SR | SS | AI | LS | WK | GT | ML | WM | LM | MM | KO | CC | JD | KL | SO | GI | SI | HM |
|---|---|---|---|---|---|---|---|---|---|---|---|---|---|---|---|---|---|---|---|---|
| CA | – | | | | | | | | | | | | | | | | | | | |
| RM | 0.028 | – | | | | | | | | | | | | | | | | | | |
| SR | 0.035 | 0.014 | – | | | | | | | | | | | | | | | | | |
| SS | 0.044 | 0.023 | 0.021 | – | | | | | | | | | | | | | | | | |
| AI | 0.063 | 0.040 | 0.038 | 0.041 | – | | | | | | | | | | | | | | | |
| LS | 0.071 | 0.046 | 0.045 | 0.049 | 0.060 | – | | | | | | | | | | | | | | |
| WK | 0.069 | 0.045 | 0.046 | 0.050 | 0.062 | 0.048 | – | | | | | | | | | | | | | |
| GT | 0.069 | 0.045 | 0.046 | 0.051 | 0.065 | 0.044 | 0.030 | – | | | | | | | | | | | | |
| ML | 0.120 | 0.108 | 0.109 | 0.110 | 0.121 | 0.010 | 0.092 | 0.078 | – | | | | | | | | | | | |
| WM | 0.121 | 0.107 | 0.108 | 0.110 | 0.120 | 0.096 | 0.089 | 0.074 | 0.081 | – | | | | | | | | | | |
| LM | 0.124 | 0.109 | 0.110 | 0.112 | 0.121 | 0.094 | 0.085 | 0.069 | 0.080 | 0.073 | – | | | | | | | | | |
| MM | 0.117 | 0.101 | 0.102 | 0.105 | 0.114 | 0.089 | 0.079 | 0.062 | 0.076 | 0.074 | 0.063 | – | | | | | | | | |
| KO | 0.121 | 0.105 | 0.106 | 0.107 | 0.118 | 0.092 | 0.080 | 0.060 | 0.076 | 0.074 | 0.067 | 0.058 | – | | | | | | | |
| CC | 0.122 | 0.107 | 0.108 | 0.110 | 0.122 | 0.097 | 0.086 | 0.068 | 0.077 | 0.078 | 0.072 | 0.064 | 0.057 | – | | | | | | |
| JD | 0.102 | 0.087 | 0.089 | 0.091 | 0.102 | 0.086 | 0.090 | 0.084 | 0.104 | 0.102 | 0.103 | 0.099 | 0.104 | 0.104 | – | | | | | |
| KL | 0.076 | 0.058 | 0.060 | 0.065 | 0.078 | 0.066 | 0.069 | 0.064 | 0.103 | 0.101 | 0.102 | 0.097 | 0.101 | 0.102 | 0.022 | – | | | | |
| SO | 0.100 | 0.083 | 0.085 | 0.088 | 0.099 | 0.080 | 0.085 | 0.078 | 0.100 | 0.098 | 0.099 | 0.094 | 0.099 | 0.099 | 0.019 | 0.027 | – | | | |
| GI | 0.106 | 0.090 | 0.091 | 0.095 | 0.104 | 0.087 | 0.092 | 0.086 | 0.104 | 0.101 | 0.102 | 0.098 | 0.102 | 0.103 | 0.015 | 0.029 | 0.015 | – | | |
| SI | 0.108 | 0.094 | 0.095 | 0.098 | 0.107 | 0.092 | 0.097 | 0.091 | 0.110 | 0.109 | 0.109 | 0.105 | 0.110 | 0.111 | 0.016 | 0.030 | 0.024 | 0.022 | – | |
| HM | 0.104 | 0.088 | 0.090 | 0.093 | 0.103 | 0.087 | 0.092 | 0.086 | 0.103 | 0.102 | 0.103 | 0.099 | 0.103 | 0.103 | 0.008 | 0.022 | 0.018 | 0.014 | 0.013 | – |

Table S4: Pairwise $F_{\mathrm{ST}}$ estimates among bighorn populations. Populations are colored by lineage: purple = Rocky Mountain; yellow = Sierra; red = desert; blue = California. Population codes: CA = Cadomin; RM = Ram Mountain; SR = Sheep River; SS = South Salmo; AI = Antelope Island; LS = Lower Salmon; WK = Whiskey Mountain; GT = Georgetown; ML = Mount Langley; WM = White Mountains; LM = Lone Mountain; MM = Muddy Mountains; KO = Kofa; CC = Carrizo Canyon; JD = Junction Deer Park; KL = Kamloops Lake; SO = South Okanagan; GI = Gilpin; SI = Sinlahekin; HM = Hart Mountain.

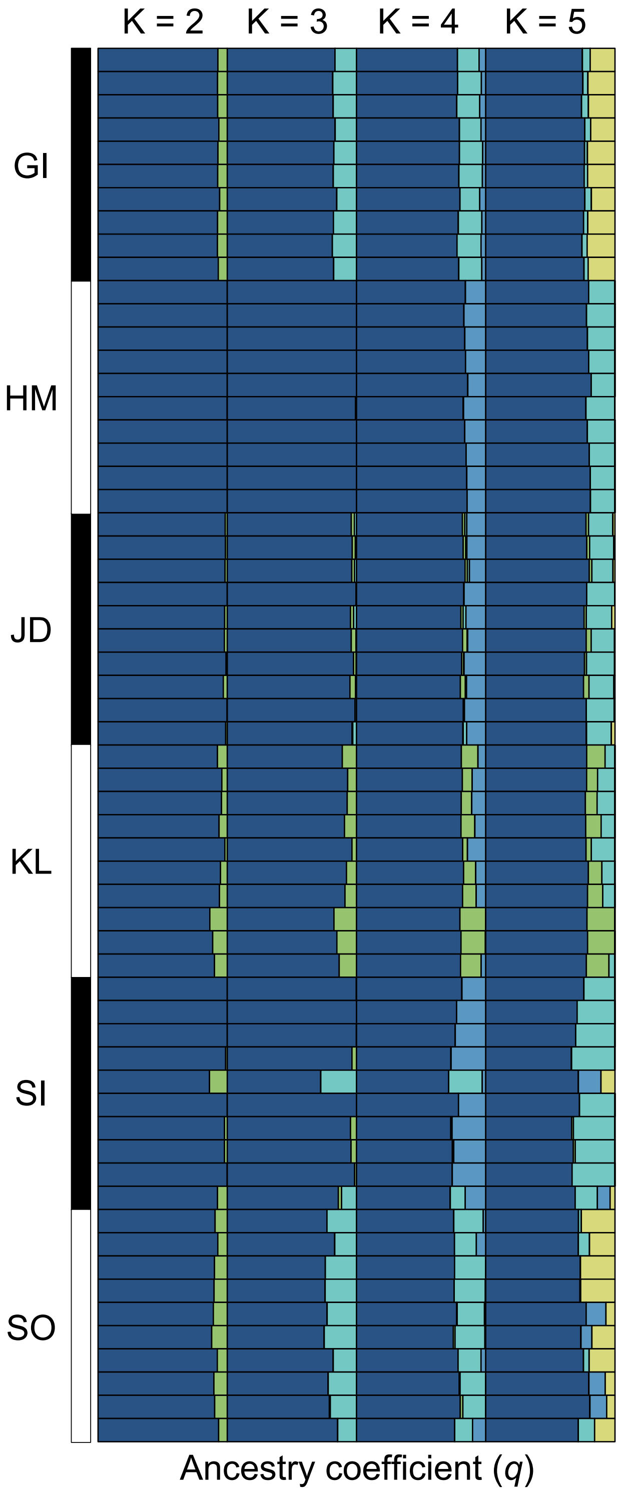

Figure S1: Individual ancestry coefficients ($q$) are shown for the California subset `entropy` models ranging from 2 – 5 genetic clusters (K). Each row represents an individual bighorn sheep, and individuals are sorted by population.

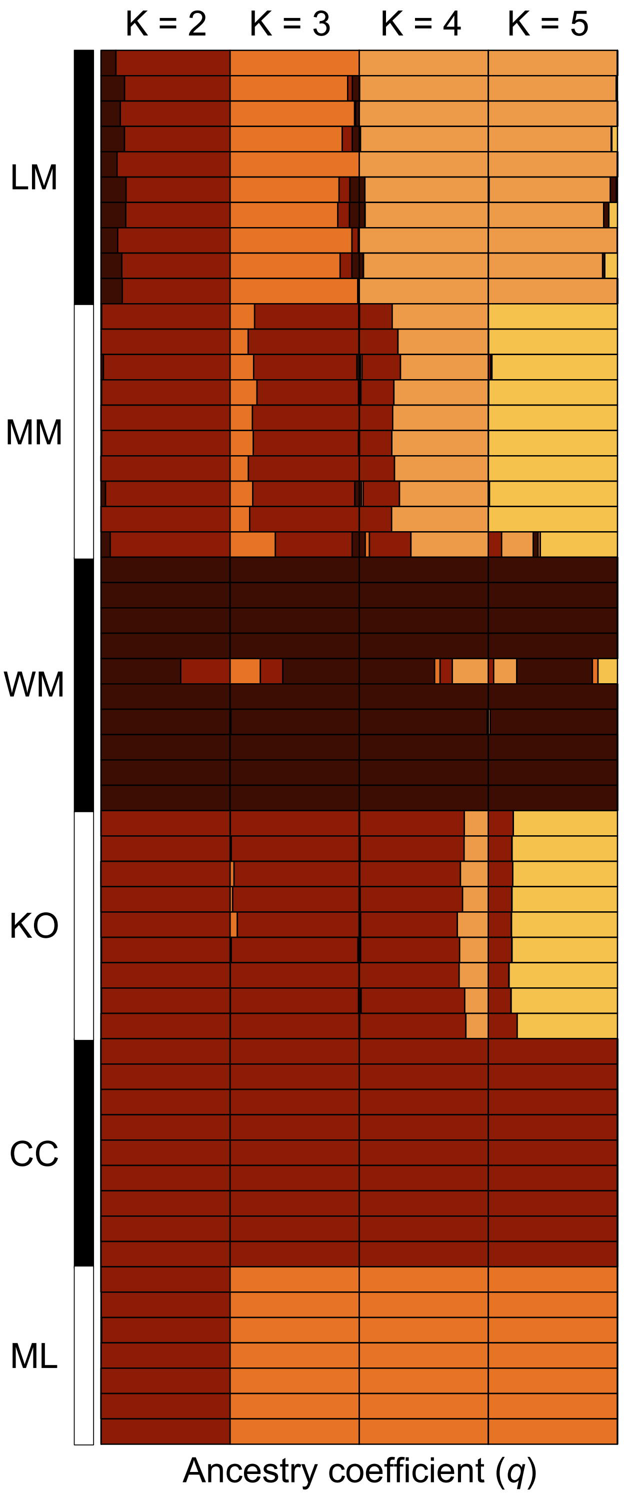

Figure S2: Individual ancestry coefficients ($q$) are shown for the desert plus Sierra subset `entropy` models ranging from 2 – 5 genetic clusters (K). Each row represents an individual bighorn sheep, and individuals are sorted by population.

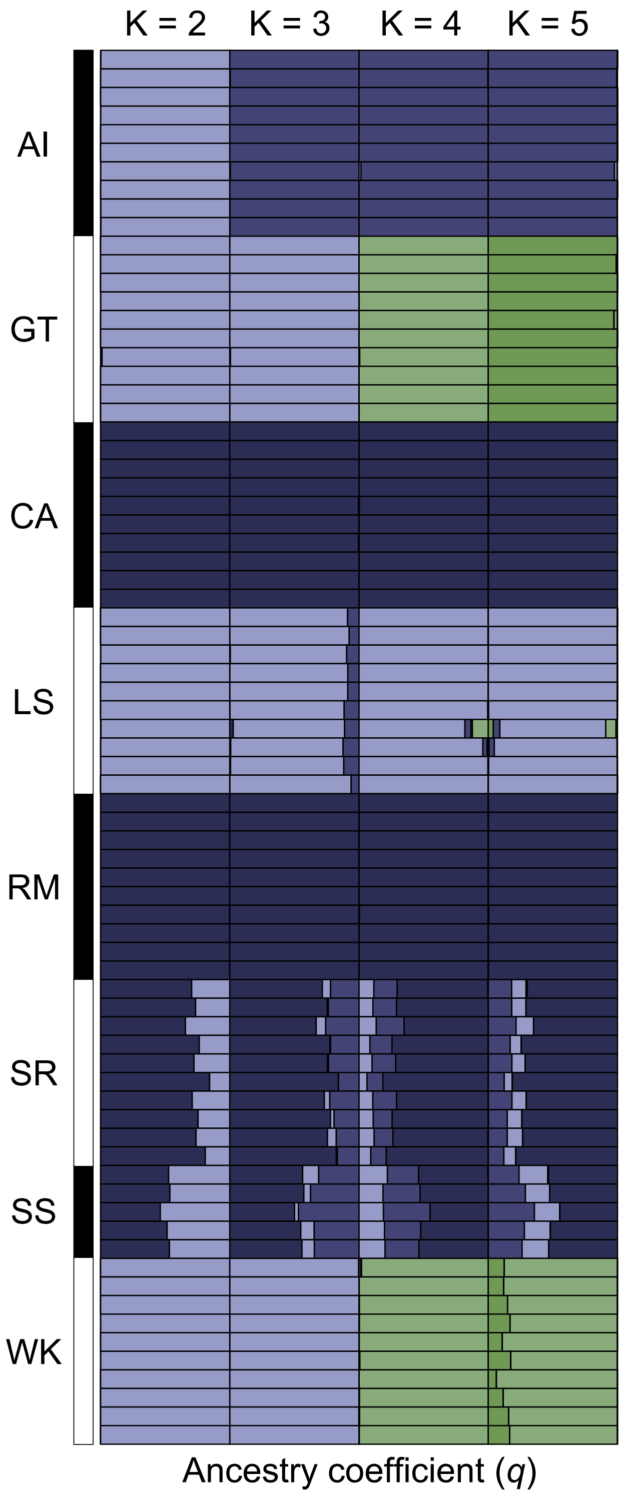

Figure S3: Individual ancestry coefficients ($q$) are shown for the Rocky Mountain subset `entropy` models ranging from 2 – 5 genetic clusters (K). Each row represents an individual bighorn sheep, and individuals are sorted by population.
